# Sensorimotor hallucinations in Parkinson’s disease

**DOI:** 10.1101/2020.05.11.054619

**Authors:** Fosco Bernasconi, Eva Blondiaux, Jevita Potheegadoo, Giedre Stripeikyte, Javier Pagonabarraga, Helena Bejr-Kasem, Michela Bassolino, Michel Akselrod, Saul Martinez-Horta, Fred Sampedro, Masayuki Hara, Judit Horvath, Matteo Franza, Stéphanie Konik, Matthieu Bereau, Joseph-André Ghika, Pierre R. Burkhard, Dimitri Van De Ville, Nathan Faivre, Giulio Rognini, Paul Krack, Jaime Kulisevsky, Olaf Blanke

**Affiliations:** Laboratory of Cognitive Neuroscience, Center for Neuroprosthetics & Brain Mind Institute, Ecole Polytechnique Fédérale de Lausanne (EPFL), Geneva, Switzerland; Movement Disorders Unit, Neurology Department Sant Pau Hospital, Barcelona, Spain; Universitat Autònoma de Barcelona (UAB), Spain; Centro de Investigación en Red-Enfermedades Neurodegenerativas (CIBERNED), Spain; Biomedical Research Institute (IIB-Sant Pau), Barcelona, Spain; University Hospital of Lausanne, CHUV, Lausanne, Switzerland; Department of Neurology, Geneva University Hospitals, Geneva, Switzerland; Department of Neurology, Besançon University Hospital, Besançon, France; Graduate School of Science and Engineering, Saitama University, Japan; Department of Neurology, Hôpital du Valais, Sion, Switzerland; Laboratoire de Psychologie et Neurocognition, LPNC, CNRS 5105 Université Grenoble Alpes, France; Medical Image Processing Laboratory, Institute of Bioengineering, Ecole Polytechnique Fédérale de Lausanne (EPFL), Lausanne, Switzerland; Department of Radiology and Medical Informatics, University of Geneva, Geneva, Switzerland; Department of Neurology, Inselspital, University Hospital and University of Bern, Bern, Switzerland

**Keywords:** Parkinson’s disease, Hallucinations, Sensorimotor, fMRI, Cognitive decline

## Abstract

Hallucinations in Parkinson’s disease (PD) are one of the most disturbing non-motor symptoms, affect half of the patients, and constitute a major risk factor for adverse clinical outcomes such as psychosis and dementia. Here we report a robotics-based approach, enabling the induction of a specific clinically-relevant hallucination (presence hallucination, PH) under controlled experimental conditions and the characterization of a PD subgroup with enhanced sensorimotor sensitivity for such robot-induced PH. Using MR-compatible robotics in healthy participants and lesion network mapping analysis in neurological non-PD patients, we identify a fronto-temporal network that was associated with PH. This common PH-network was selectively disrupted in a new and independent sample of PD patients and predicted the presence of symptomatic PH. These robotics-neuroimaging findings determine the behavioral and neural mechanisms of PH and reveal pathological cortical sensorimotor processes of PH in PD, identifying a more severe form of PD associated with psychosis and cognitive decline.

## Introduction

The vivid sensation that somebody is nearby when no one is actually present and can neither be seen nor heard (i.e. sense of presence or presence hallucination, PH), has been reported from time immemorial and found its way into the language and folklore of virtually all cultures^1–3^. Following anecdotal reports of PH by extreme mountaineers^4^, solo-sailors and shipwreck survivors^5^, PH have also been described in a variety of medical conditions including schizophrenia^1,6^, epilepsy, stroke, brain tumors^7–9^ and Parkinson’s disease (PD)^10–12^.

Whereas PH are rare manifestations in most medical conditions, they are frequent and may occur regularly, even on a daily basis, in patients with PD. Hallucinations, including PH, are not only frequent, occurring in up to 60% of PD patients, but increase in frequency and severity with disease progression and are one of the most disturbing non-motor symptoms^11–13^. Importantly, PH and other hallucinations in PD are associated with major negative clinical outcomes such as chronic psychosis, cognitive decline and dementia, as well as higher mortality^10,11,14–16^. PH are generally grouped with so-called minor hallucinations and are the most prevalent and earliest type of hallucination in PD^11,12^, often preceding the onset of structured visual hallucinations^17^, and may even be experienced, by one-third of patients, before the onset of first motor symptoms^18^. Despite their high prevalence and strong association with major negative clinical outcome, PH (and other hallucinations) remain underdiagnosed^12,14,19,20^, caused by patients’ reluctance to report hallucinations and difficulties to diagnose and classify them^21,22^.

Past research described changes in visual function, cognitive deficits and related brain mechanisms in PD patients with hallucinations, yet these studies focused on patients with structured visual hallucinations^23^. Comparable studies are rare or lacking for PH (or other minor hallucinations) and very little is known about the early brain dysfunction of PH in PD and how they lead to more severe and disabling structured visual hallucinations and cognitive deficits^11,24^. Early neurological work investigated PH following focal brain damage and classified PH among disorders of the body schema, suggesting that they are caused by abnormal self-related bodily processes^9,25^.

More recent data corroborated these early findings and induced PH repeatedly by electrical stimulation of a cortical region involved in sensorimotor processing^8^. By integrating these clinical observations with human neuroscience methods inducing bodily illusions^27–30^, we have designed a method able to robotically induce PH (robot-induced PH or riPH) in healthy participants^26^. This research demonstrated that specific sensorimotor conflicts, including bodily signals from the arm and trunk, are sufficient to induce mild to moderate PH in healthy participants, linking PH to the misperception of the source and identity of sensorimotor signals of one’s own body.

Here, we adapted our robotic procedure to PD patients and elicited riPH, allowing us to characterize a subgroup of patients that is highly sensitive to the sensorimotor procedure, and to identify their aberrant sensorimotor processes (study 1). We next determined the common PH-network in frontal and temporal cortex, by combining MR-compatible robotics in healthy participants with brain network analysis in neurological non-PD patients with PH (study 2). Finally, we recorded resting-state fMRI data in a new and independent sample of PD patients and identified pathological functional connectivity patterns within the common PH-network, which were predictive for the occurrence of PD-related PH (study 3).

## Results

### riPH in patients with PD (study1.1)

Based on semi-structured interviews, patients with PD were grouped into those who reported symptomatic PH, sPH (PD-PH; n=13), and those without sPH (PD-nPH; n=13) (Supplementary S1-2, Tab.S1-2). Patients were asked to actuate a robotic device and were exposed to repetitive sensorimotor stimulation that has been shown to induce PH in healthy participants in a controlled way^26^. In study1.1, we assessed whether robotic sensorimotor stimulation induces PH in patients with PD and whether such riPH differ between PD-PH and PD-nPH, hypothesizing that PD-PH patients are more sensitive to the robotic procedure.

In the robotic sensorimotor paradigm, participants were asked to perform repetitive movements to operate a robot placed in front of them, which was combined with a back robot providing tactile feedback to participants’ backs (Fig.1A). Based on previous data^26,28,31^, tactile feedback was delivered either synchronously with patients’ movements (synchronous control condition, a spatial conflict is present between movement in front and touch on the back) or with a 500ms delay (asynchronous condition) associated with an additional spatio-temporal sensorimotor conflict shown previously to induce PH^26,36^ (Supplementary S3).

**Figure 1.**
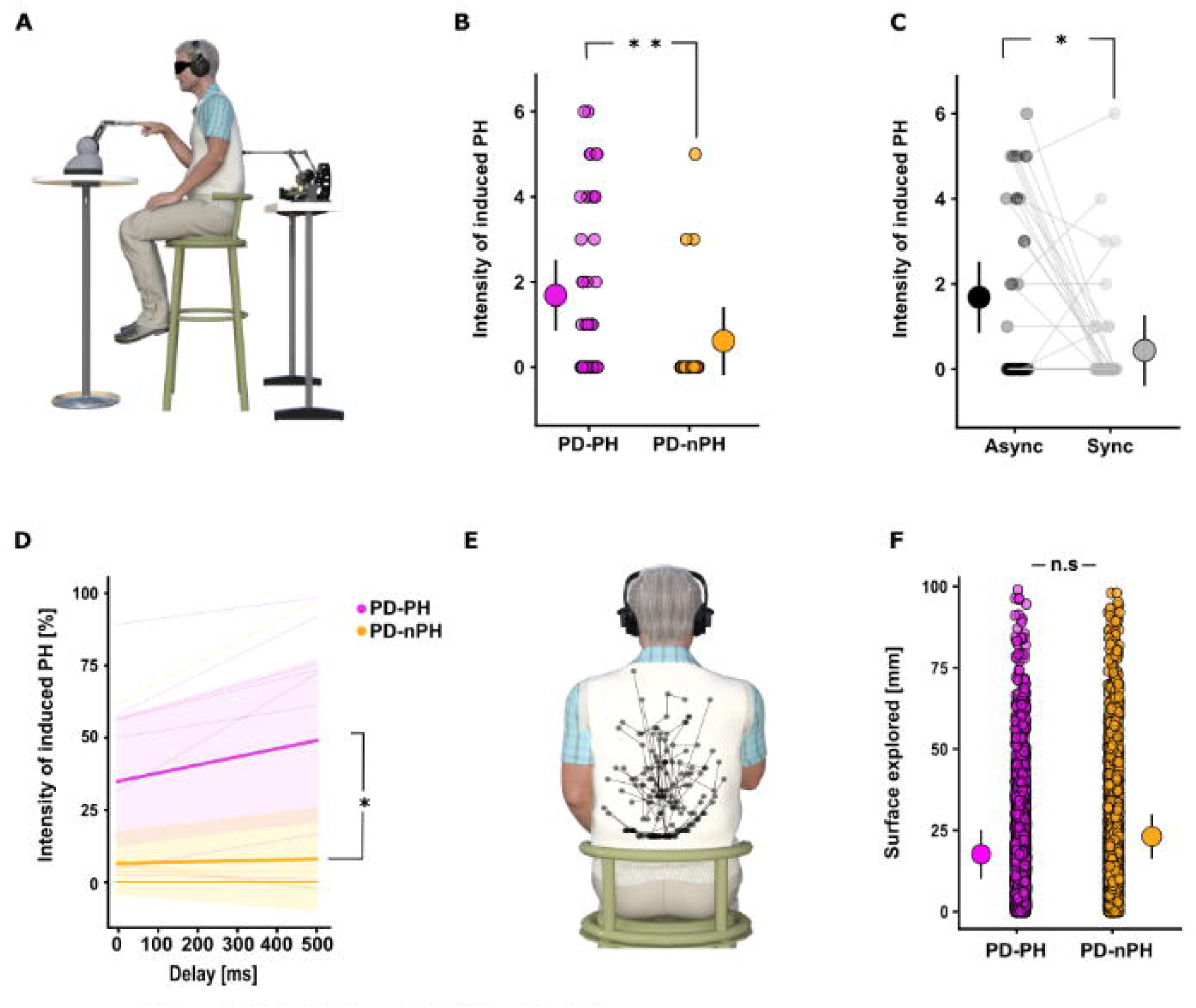
Robot-induced PH (PD patients). **A.** Setup for study 1. Responses in synchronous and asynchronous conditions are shown. During the asynchronous condition, the sensorimotor feedback on the participants’ back was delayed by 500 ms (study1.1) or with a random delay (0-500ms, steps of 100ms) (study1.2). **B.** Study1.1. riPH in PD-PH are stronger than in PD-nPH. Each dot indicates the individual rating of the intensity of the riPH (PD-PH (purple) and PD-nPH (yellow)). The dot with the bar on the left and right side indicate the mixed effects linear regression between PD-PH and PD-nPH. Error bar represent 95% confidence interval. **C.** Study1.1. Asynchronous condition induced stronger riPH. Each dot indicates the individual rating of the intensity of the riPH. The dot with the bar on the left and right side indicate the mixed effects linear regression between Asynchronous (black) and Synchronous (gray) sensorimotor stimulation. Error bars represent 95% confidence interval. **D.** Study1.2. riPH were modulated by delay (permutation p-value=0.014) and PD-PH vs. PD-nPH were more sensitive to the sensorimotor stimulation (slope permutation p-value=0.039, intercept p-value=0.016). The thicker line indicates the mean of the fitted models, the shaded are indicates the 95% confidence interval, thinner lines indicate single subject fit. **E.** Study1.2. Exemplary movements executed by one patient during sensorimotor stimulation. **F.** Study1.2. Mixed effects linear regression between the Euclidean distance between pokes for PD-PH (purple) and PD-nPH (yellow). Error bar represent 95% confidence interval.

The robotic procedure was able to induced PHs in patients with PD. Importantly, PD-PH patients rated the intensity of riPH higher than PD-nPH patients (main effect of Group: permutation p-value=0.01) (Fig.1B). Confirming the general importance of conflicting asynchronous sensorimotor stimulation^26^ for riPH, both sub-groups gave higher PH ratings in the asynchronous versus synchronous condition (main effect of Synchrony: permutation p-value=0.045) (Fig.1C) (Supplementary S4 for additional results). Other robot-induced bodily experiences (e.g. illusory self-touch) also confirmed previous findings^26^ (Supplementary S5) and no differences were observed for the control items (all permutation p-values>0.05). These results show that PH can be safely induced by the present robotic procedure under controlled conditions in patients with PD. Such riPH were modulated by sensorimotor stimulation with asynchronous robotic stimulation resulting in higher ratings in all tested groups, and, importantly, PD-PH (vs. PD-nPH) reported stronger riPH, linking the patients’ usual sPH to experimental riPH and showing that PD-PH patients were more sensitive to our robotic procedure.

Post-experiment debriefing revealed 38% of PD-PH patients who reported riPH that were comparable (or even stronger) in intensity to the patients’ usual sPH in daily life. One PD-PH patient, for example, described his riPH as “an adrenaline rush. Like something or someone was behind me, although there is no possibility to have someone behind” (for additional reports Supplementary S6). Interestingly, all such instances were reported after asynchronous stimulation. Moreover, PD-PH patients often experienced riPH on their side (and not on their back, where tactile feedback was applied), revealing a further phenomenological similarity between riPH and PD patients’ usual sPH^10^ and suggesting that we induced a mental state that mimics sPH (Supplementary S7-8).

Data from study1.1 reveal that riPH can be safely induced by the present procedure, are stronger in patients who report sPH (PD-PH), and that such riPH share phenomenological similarities with PD-related sPH. These findings cannot be related to a general response bias related to PD, because riPH were absent or weaker in PD-nPH and because the control items showed no effects in any of the participant groups.

### riPH in PD-PH patients depend on sensorimotor delay (study1.2)

Previous work investigated the effects of systematically varied sensorimotor conflicts (i.e. delays) on somatosensory perception, enabling the induction and modulation of different somatic experiences and illusions^31–33^. Sensorimotor processing and the forward model of motor control^34,35^ are prominent models of hallucinations^36,37^ and it has been proposed that deficits in predicting sensory consequences of actions causes abnormal perceptions and hallucinations^36–38^. In study1.2, we assessed whether riPH depend on the degree of conflict applied during sensorimotor stimulation, by inserting variable delays between the movements of the front robot (capturing movements of the forward-extended arm) and the back robot (time of tactile feedback on the back). In each trial, participants (Supplementary S9) were exposed to a randomly chosen delay (0-500ms, steps of 100ms). After each trial, participants were prompted whether they experienced a riPH or not (yes-no response, Supplementary S10). We investigated whether the intensity of riPH increases with increasing delays in PD patients (showing that PH are modulated by increasing spatio-temporal conflicts) and whether PD-PH have a higher spatio-temporal delay sensitivity than PD-nPH.

As predicted, study1.2 shows that the intensity of riPH increased with increasing spatio-temporal conflict (main effect of delay: permutation p-value=0.014) and that this delay dependency differed between the two patient groups, showing a higher delay sensitivity in PD-PH patients (interaction Group*delay: permutation p-value=0.039) (Fig.1D) (Supplementary S11, Fig.S1). Control analysis (Supplementary S12) (Fig.1E-F, Fig.S2) allowed us to exclude that the observed differences (in riPH ratings between patient groups) are due to differences in movements of the arm and related tactile feedback during the robot actuation (Supplementary S13). In addition, these differences in riPH between PD-PH and PD-nPH cannot be explained by differences in demographic or clinical variables (including anti-parkinsonian medication, motor impairment; all permutation p-values>0.05) (Supplementary S14, Tab.S1).

Based on previous results using robotics and conflicting sensorimotor stimulation to alter somatosensory perception^31–33^, these data extend those of study1.1 and reveal abnormal perceptual processes in PD-PH patients when exposed to different sensorimotor conflicts, characterized by experiencing stronger riPH and a higher sensorimotor sensitivity. These findings are compatible with an alteration of sensorimotor brain processes associated with the forward model and its role in hallucinations in PD-PH patients^36,37,39^.

### Brain mechanisms of PH

Neuroimaging work on sPH and other minor hallucinations in PD patients has described structural alterations and aberrant functional connectivity in different cortical regions^24,40^. Despite these clinical neuroimaging findings, it is not known whether the regions associated with sPH of neurological non-parkinsonian origin^26^ are also altered in PD patients with PH. Moreover, because the brain networks of riPH have never been investigated, it is also not known whether the abnormal sensorimotor mechanisms described in PD-PH patients (study1) are associated with a disruption of brain networks of riPH. To determine the brain mechanisms of PH, we first adapted an MR-compatible robot^41^ (Supplementary S15) and applied sensorimotor stimulations while recording fMRI during riPH in healthy participants and identified the associated brain networks (study2.1). We then combined this network with evidence from sPH of neurological non-parkinsonian origin (study 2.2) and, finally, applied this common network to PD patients (study 3).

### Brain mechanisms of riPH in healthy participants using MR-compatible robotics (study2.1)

Based on behavioral pilot data (Supplementary S16-S17, Tab.S5), we exposed 25 healthy participants to asynchronous and synchronous robotic stimulation while recording fMRI (Fig.2A, Supplemental S15, Fig.S3). Our behavioral data replicated previous results (^26^, study1 and pilot study) and we found that asynchronous vs. synchronous robotic stimulation induces stronger PH (main effect of Synchrony: permutation p-value=0.0082, Fig.2B) and another bodily experience (Tab.S6), but did not modulate control items (all permutation p-values>0.08, Supplementary S18, Tab.S6). As for study1.2, riPH were not related to movement differences across conditions (permutation p-value=0.99) (Fig.2C), confirming that sensorimotor stimulation (and not movement differences) applied with the MR-compatible robot modulated PH intensity across conditions.

**Figure 2.**
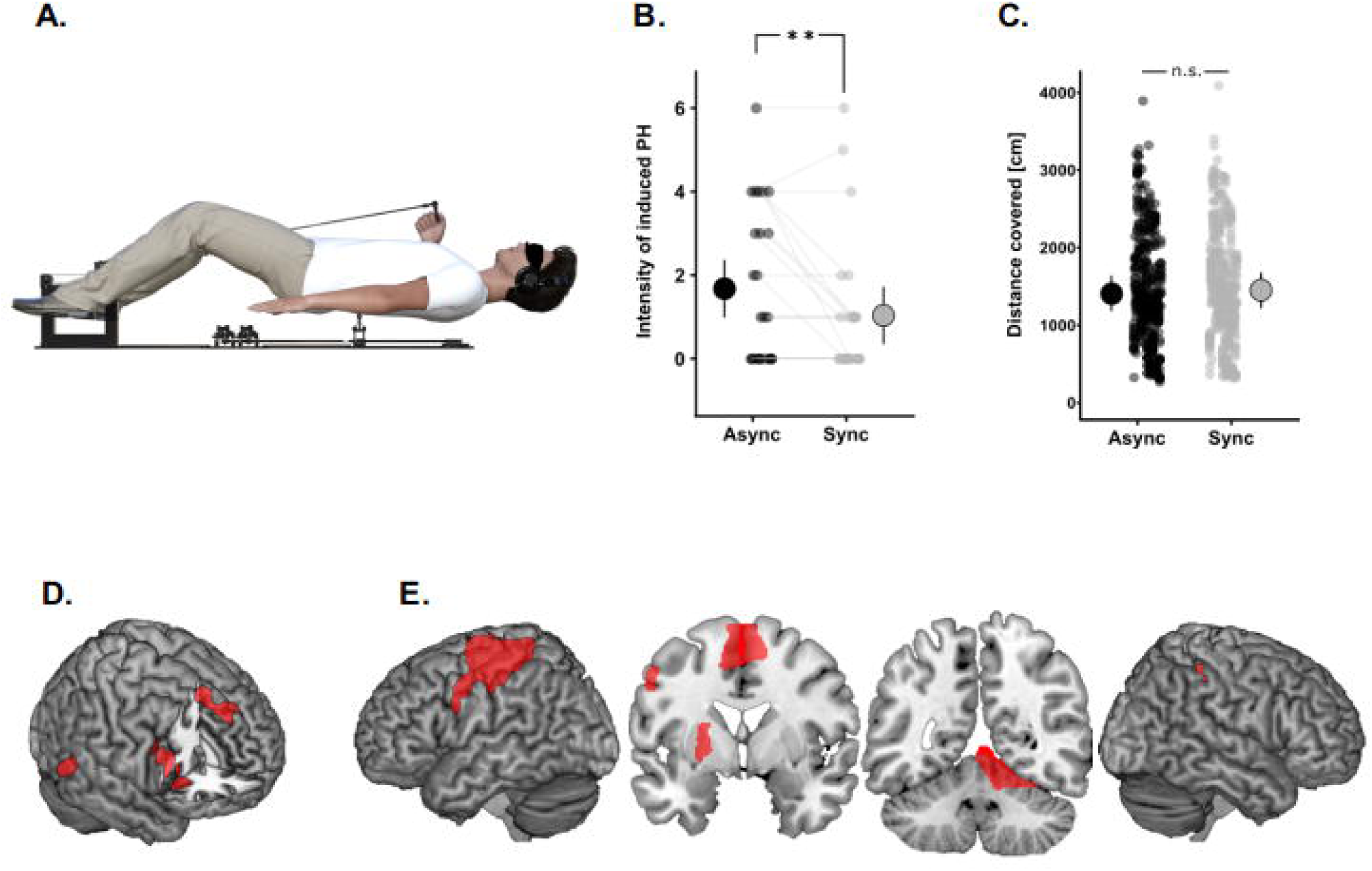
Neuroimaging results of robot-induced PH (healthy participants). **A.** MR-compatible robotic system is shown. Participants were instructed to move the front robot with their right hand and the back robot delivered the touch to the participant’s back either synchronously or asynchronous (500ms delay between their movement and the sensory feedback received on the back). **B.** Asynchronous vs. synchronous condition induced stronger riPH. Each dot indicates the individual rating of the intensity of the riPH in healthy participants. The dot with the bar on the left and right side indicate the mixed effects linear regression between asynchronous (black) and synchronous (gray) sensorimotor stimulation. Error bar represents 95% confidence interval. **C.** Movement data from the fMRI experiment: no movement differences were found between the two conditions. **D.** Brain regions sensitive to the delay. **E.** Brain areas present in the conjunction analysis between the contrast synchronous>motor+touch and the contrast asynchronous>motor+touch. The coronal slices are at Y = −1 and Y = −53. There was no anatomical overlap between both networks (D and E).

To identify the neural mechanisms of riPH, we determined brain regions that were (1) more activated during the asynchronous vs. synchronous condition (spatio-temporal sensorimotor conflict) and (2) activated by either of the sensorimotor conditions (synchronous, asynchronous) vs. two control conditions (motor and touch) (Supplementary S19, conjunction analysis). Regions more activated during asynchronous vs. synchronous sensorimotor stimulation were restricted to cortical regions (Fig.2D, Tab.S7) and included the inferior frontal gyrus (IFG), anterior insula, medial prefrontal cortex (mPFC) and the posterior part of the middle temporal gyrus (pMTG, bordering on angular gyrus and adjacent occipital cortex). Conjunction analysis (between contrast synchronous>motor+touch and contrast asynchronous>motor+touch) (Supplementary S20, Fig.S4) revealed a subcortical-cortical network in left sensorimotor cortex (contralateral to the hand moving the robot, including M1, S1 and adjacent parts of premotor cortex and superior parietal lobule), in bilateral supplementary motor area (SMA), right inferior parietal cortex, left putamen, and right cerebellum (Fig.2E, Tab.S8).

Collectively, these fMRI results constitute the first delineation of the neural underpinnings of riPH in healthy participants that is unrelated to movement differences across conditions and distinct from activations in two control conditions, revealing a network of brain regions that have been shown to be involved in sensorimotor processing and in agency (such as M1-S1, pMTG^42,43^, PMC^44,45^, SMA^43,46^, IPS^47,48^, as well as the cerebellum ^42,49^ and putamen).

### Common PH-network for sPH and riPH (study2.2)

To determine neural similarities between riPH and sPH and confirm the sensorimotor contribution to sPH, we first applied lesion network mapping (Supplementary S21) and identified network connectivity mapping in neurological non-parkinsonian patients, in whom sPH were caused by focal brain damage (study2.2), and then determined the common network (cPH-network) between the riPH and sPH. Lesion network mapping^50^ extends classical lesion symptom mapping by considering each lesion as a seed (region of interest, ROI) and computing its connectivity map (in normative resting state fMRI data, publicly available database, 126 healthy participants^51^) (Fig.S5).

This analysis revealed that all lesions had functional connectivity with bilateral posterior superior temporal gyrus/temporo-parietal junction (pSTG/TPJ), bilateral middle cingulate cortex (MCC), bilateral insula, and right IFG, constituting the sPH-network (Fig.3A, for all regions see Tab.S9) and did not overlap with connectivity patterns of a control hallucination network (Supplementary S22-S23, Tab.S10). We then determined the common regions between the sPH-network (non-parkinsonian neurological patients) and the riPH network (healthy participants). This cPH-network consisted of three regions, including right IFG, right pMTG, and left vPMC (Fig.3B, Supplementary S24) and is the first neuroimaging evidence that riPH and sPH recruit similar brain regions, even if both types of PH differ in several aspects such as frequency, intensity, trigger mechanism, supporting a link between sensorimotor robotics inducing hallucinatory states with neuroimaging in healthy participants and in patients.

**Figure 3.**
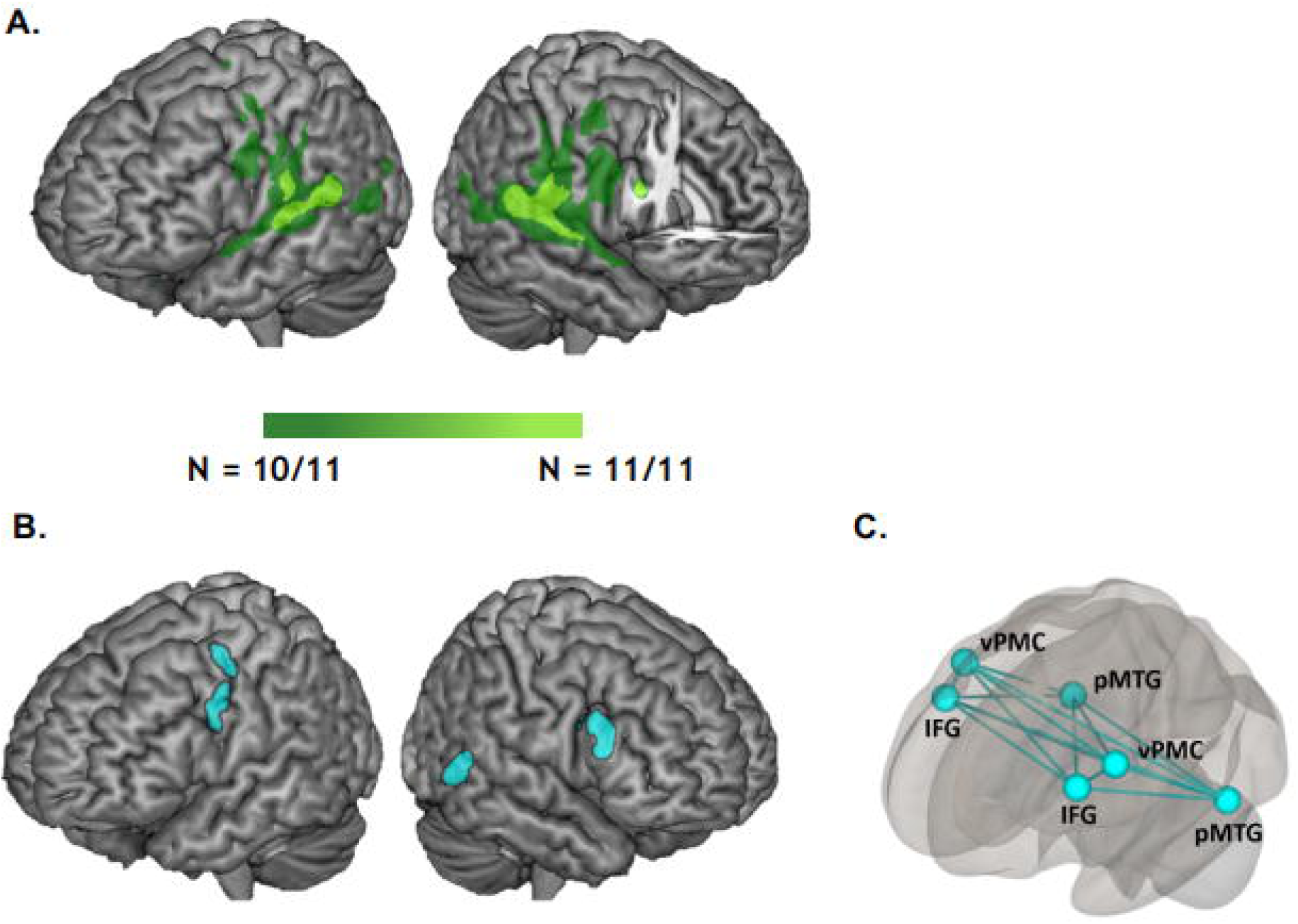
Symptomatic PH-network and common PH-network. **A.** sPH network connectivity in neurological non-parkinsonian patients. **B.** Common regions between the riPH-network and sPH-network (cPH-network) were found in three regions: left vPMC, right IFG and right pMTG. **C.** Schematic display of the cPH-network projected bilaterally.

### Disrupted functional connectivity in cPH-network accounts for sPH Parkinson’s disease (study3.1)

To assess the relevance of the cPH-network for PD patients’ usual sPH in daily life, we analyzed resting state fMRI data in a new group of PD patients and investigated whether functional connectivity of the cPH-network (as defined in study2, projected bilaterally, Fig.3C) differed between PD-PH and PD-nPH (new cohort of 30 PD patients) (Supplementary S25-26, Tab.S11). Based on the disconnection hypothesis of hallucinations^52^, evidence of decreased connectivity for hallucinations of psychiatric origin^37^, and aberrant functional connectivity in PD patients with minor hallucinations including PH^24^, we predicted that the functional connectivity within the cPH-network differs between both PD patient groups and that the connectivity within the cPH-network is reduced in PD-PH vs. PD-nPH patients. We found that the functional connectivity within the cPH-network, predicted with 93.7% accuracy whether a patient was clinically classified as PD-PH (kappa:0.86, permutation p-value=0.0042). Moreover, within the cPH-network, the functional connectivity between the left IFG and left pMTG contributed mostly to the classification of the two sub-groups (Tab.S12). PD-PH had reduced IFG-pMTG connectivity (permutation p-value<0.0001; Fig.4A-B). These changes were selective because (1) the same analysis in a control network (Fig.S7) (same size, same number of connections) did not predict the occurrence of hallucinations based on the functional connectivity (accuracy:27.7%, kappa:-0.43, permutation p-value=0.24) and (2) no changes in functional connectivity were observed when analyzing whole brain connectivity. These data show that reduced fronto-temporal connectivity within the cPH-network distinguishes PD patients with sPH from those without hallucinations, in accordance with the disconnection hypothesis of hallucinations^52–54^.

**Figure 4.**
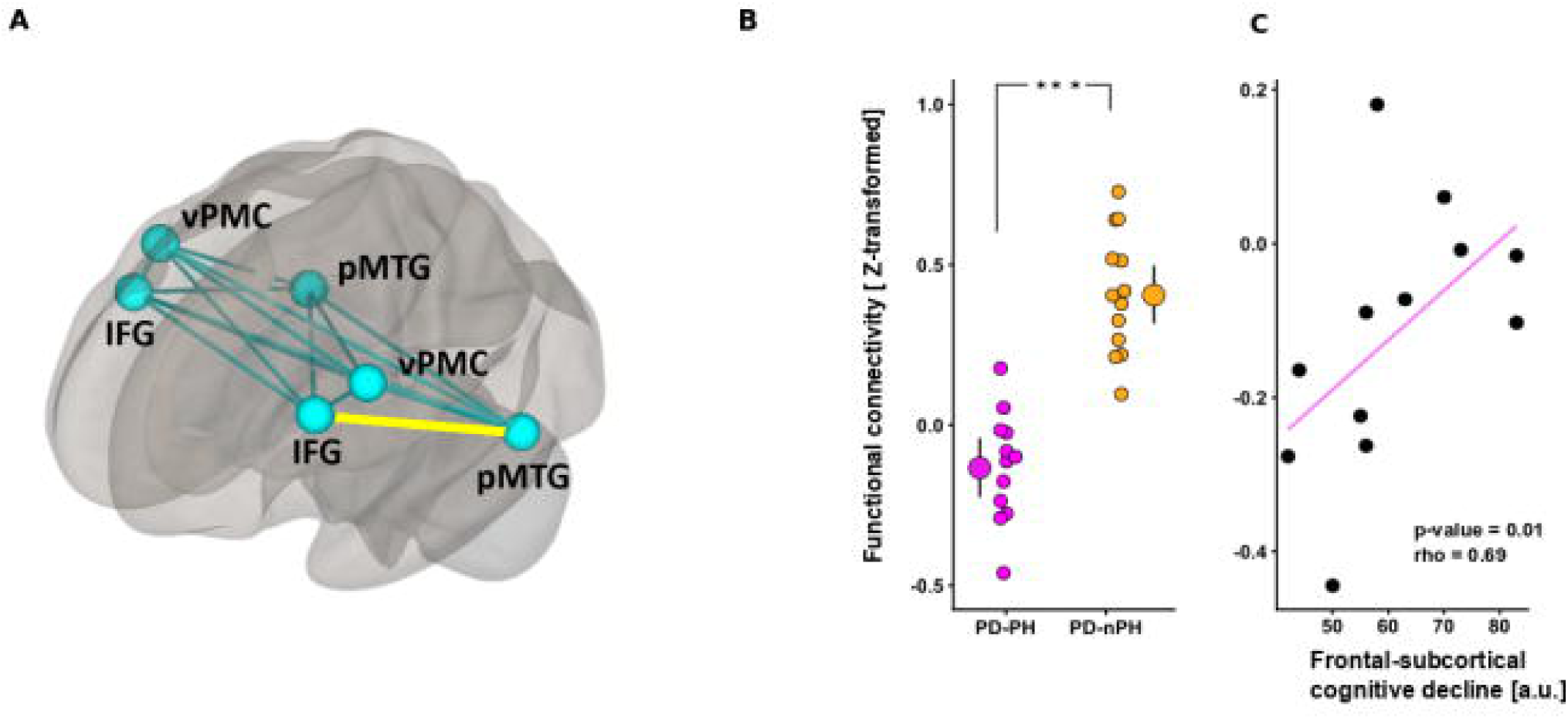
Functional connectivity in the sensorimotor network. **A.** Connections showing differences in functional connectivity between PD-PH vs. PD-nPH within the cPH-network are shown (yellow). **B.** Mixed effects linear regression between the functional connectivity for PD-PH (purple) and PD-nPH (yellow) between left IFG and left pMTG is shown. PD-PH vs. PD-nPH patients have a significantly reduced functional connectivity. Error bar represents 95% confidence interval, and the dot represents the mean functional connectivity. Dots represent the functional connectivity for each patient. **C.** Degree of functional disconnection is correlated with the cognitive decline (fronto-cortical sub-score of PD-CRS) in PD-PH patients. Lower connectivity was correlated with lower frontal cognitive fronto-subcortical abilities.

### Functional disconnection within the cPH-network correlates with cognitive decline for PD-PH (study3.2)

It has been suggested that PH (and minor hallucinations) are indicative of a more severe and rapidly advancing form of PD, evolving towards structured visual hallucinations and psychosis^11,17^, as well as faster cognitive deterioration including dementia^16,55–57^. We therefore tested whether functional connectivity between the left IFG and the left pMTG within the cPH-network relates to cognitive dysfunction in the present PD-PH patients. Results show that stronger decreases in left IFG-pMTG connectivity are associated with stronger cognitive decline (PD-CRS^58^), reflecting differences in frontal-subcortical function (p-value=0.01,rho=0.69,Fig.4C), but not on posterior-cortical function (p-value=0.66, rho=-0.15, the two correlations differed significantly: t=3.87, p-value<0.01). These results reveal an association between fronto-subcortical cognitive alterations and specific decreases in fronto-temporal connectivity within the cPH-network in PD-PH patients, compatible with a more severe form of PD associating PH and cognitive decline.

### General Discussion

Having developed a robotic procedure that can induce PH in PD patients under safe and controlled sensorimotor conditions, we report that PD patients with sPH are highly sensitive to the procedure and reveal abnormal sensorimotor mechanisms leading to PH. Using MR-compatible robotics in healthy participants combined with lesion network mapping analysis in patients with sPH of neurological non-parkinsonian origin, we identify the common network associated with PH and show that fronto-temporal connectivity within this cPH-network is selectively disrupted in a new and independent sample of PD patients. Disruption of the cPH-network was only found in PD patients suffering from sPH (PD-PH) and the degree of this disruption further predicted the severity of cognitive decline.

The present behavioural findings show that stronger sensorimotor conflicts result in stronger riPH, supporting and extending previous evidence in favor of an alteration of self-related sensorimotor processing as a fundamental mechanism underlying PH^33^. Importantly, we show that this mechanism is especially vulnerable in PD-PH patients, revealed by their stronger bias and sensitivity when exposed to conflicting sensorimotor stimulation. These results extend the sensorimotor forward model to hallucinations in PD-PH patients^36,37,39^ and support earlier evidence in neurological non-PD patients that PH are self-related body schema disorders associated with altered sensorimotor self-monitoring^7–9^.

By including fMRI data from healthy participants experiencing riPH and from non-parkinsonian neurological patients with sPH, we mapped common brain structures between both types of PH, which we showed to be selectively disrupted in PD patients with sPH. The imaging results within this cPH-network further revealed aberrant functional connectivity decreases between fronto-temporal regions that have been associated with outcome processing of sensorimotor signals and the forward model^54,59^, further linking PH in PD to the fronto-temporal hallucination disconnection model^52,54,60^. The present account - involving sensorimotor mechanisms and brain structures in fronto-temporal cortex rather than posterior brain functions and regions - is functionally and conceptually distinct from earlier proposals that hallucination in PD are caused by visuo-spatial deficits^23^ or that sPH are caused by abnormal social-cognitive brain mechanisms^10^ in parietal or occipital cortex^23,61,62^. Our finding that the decreased fronto-temporal connectivity within the cPH-network is associated with stronger cognitive decline of PD-PH patients in fronto-subcortical (but not posterior-cortical, functions) lends support to clinical suggestions about the importance of PH (and other minor hallucinations) as a major risk factor not only for the occurrence of structured visual hallucinations and psychosis^17^, but also for a more severe and rapidly advancing form of PD ^11,16,55,57^.

Because the phenomenology of riPH resembles those of sPH and PD-PH patients were found to be more sensitive to riPH, the present procedure provides researchers and clinicians with new objective possibilities to assess the occurrence and intensity of subjective hallucinatory phenomena by quantifying delay-sensitivity and the repeated online induction of hallucinatory states across controlled conditions in PD patients, as well as the association of these measures with cPH-network activity. This is not possible in current clinical practice that is based on clinically important, but post-hoc interviews between physician and patient, often about hallucinations that have occurred many days or weeks ago, and that many patients hesitate to speak about^21^. The detection of specific behavioural and imaging changes associated with specific hallucinatory states that are observed online during the robotic procedure will improve the quantification and prediction of a patient’s proneness for hallucinations and psychosis and may facilitate targeted pharmacological interventions that limit side effects^63^.

## Methods

### Study 1

#### Participants (study1.1-1.2)

All participants provided written informed consent prior to the experiments. The study was approved by the Cantonal Ethics Committee of Geneva (Commission Cantonale d’Ethique de la Recherche sur l’Être Humain), the Cantonal Ethics Committee of Vaud. Participants of study1 consisted of patients with PD (n=26) and age-matched healthy controls (HC, n=21) (Supplementary S1-S4). Based on an extensive semi-structured interview (conducted after the experimental sessions) about hallucinations (including sPH), PD patients were separated into two sub-groups: patients who reported sPH as part of their PD (PD-PH) (n=13) and PD patients without sPH (PD-nPH) (n=13). Patients were considered as having sPH if they answered affirmatively to the question that previous investigators have used: “do you sometimes feel the presence of somebody close by when no-one is there?” The hallucinated presence could be located behind, on the side (left or right) of the patient, or in another room and was generally not seen (see ^2,7,8,10,26^). All PD patients, who were included in study1 presented idiopathic PD diagnosed by trained neurologists. No patient was suffering from a neurological disorder other than PD (more details in Supplementary S2).

#### General experimental procedure (study1)

Each PD patient underwent study1 at a similar time (10am), after having received their usual anti-parkinsonian medication and were in their “best ON” state. To investigate riPH, we adapted the experimental method and device as our previous research^26^. Briefly, sensorimotor stimulation was administered with a robotic system consisting of two robotic components (front-robot, back-robot) that has previously been used to induce PH. For each experimental session, we applied the following conditions: synchronous sensorimotor stimulation (the participants were asked to move the front-robot via either their right or left hand that was actuating the movements of the back-robot to apply tactile feedback to their back); asynchronous sensorimotor stimulation (same as synchronous stimulation, but with an additional temporal delay between the front-robot and the back-robot; see below for details of each experiment; Fig.1A). During sensorimotor stimulation, participants were always asked to keep their eyes closed and were exposed to continuous white noise through headphones (Supplementary S3).

#### Procedure, design, and analysis (study1.1)

Participants were asked to insert their index finger in the haptic front-robot and carry out repeated poking movements while they received tactile cues on their backs, delivered by the back-robot. Thus, sensorimotor stimulation included motor, tactile, and proprioceptive signals from the upper limb moving the front-robot and additional tactile signals from the back-robot. Stroking was applied either synchronously (0ms delay) or asynchronously (500ms delay) (*Synchrony*: asynchronous vs. synchronous). Additionally, we measured the effect of the side of the body (i.e. hand moving the front-robot) that was most strongly affected by PD versus the other hand (*Side*) to investigate if the hemisphere predominantly affected by PD influenced riPH^64,65^. The factors (*Synchrony*; *Side*) and the order of testing were randomized across participants. Each participant randomly started with one Side first, for which the two *Synchrony* conditions (random order) were tested, and then the second Side was tested with the two *Synchrony* conditions (random order). In total, each participant performed four sessions (one per condition) lasting two minutes each. At the end of each of the four sensorimotor stimulation conditions, all participants filled a questionnaire (see below). Each PD-PH, PD-nPH, and HC included in the study was able to perform the entire study1.1.

##### PH and other subjective ratings

To measure PH and other illusions, we administered a questionnaire (6 questions) that was adapted from^26^. Participants were asked to indicate on a 7-point Likert scale, how strongly they felt the sensation described by each item (from 0 = not at all, to 6 = very strong). For questions see Supplementary S5.

##### Data analysis

Each question was analyzed with linear mixed effects models (lme4 and lmerTest both R packages^66,67^). Models were performed on the subjective ratings in each of the four conditions with *Synchrony* (synchronous vs. asynchronous), *Groups* (i.e., PD-PH vs. PD-nPH, and PD-PH vs. HC) and *Side* as fixed effects, and random intercepts for each subject. The significance of fixed effects was estimated with a permutation test (5000 iterations; predictmeans^68^ R package).

#### Procedure, design, and analysis (study1.2)

To complement and extend study1.1, we applied a Yes/No task, following sensorimotor stimulation, in which participants were asked to report whether they experienced a PH or not, on a trial-by-trial basis. On each sensorimotor stimulation trial, the delay between the movement and the stroking on the back was randomly chosen from a delay between 0 and 500ms (steps of 100m). One trial started with an acoustic signal (400 Hz tone, 100ms duration) indicating the beginning of the trial: at this point the participant started with the poking movements. Once the number of pokes reached a total of six (automatically counted), two consecutive tones (400 Hz, 100ms duration) indicated to the participant to stop the movements and to verbally answer with either a “Yes” or a “No” to the PH question, (Question: “Did you feel as if someone was standing close by (behind you or on one side)?”). The investigators where always placed > 4 meters away and in front from the participants during the experiment. Each participant was asked to perform three sessions; each session consisted of 18 trials (3 repetitions per delay (9 repetitions in total)). Between each session, the participant could take a break according to his/her needs (Supplementary S10).

##### riPH rating analysis

First, to investigate how the degree of sensorimotor conflict modulates PH, we analyzed the behavioral responses as a function of different delays (i.e., 0-500ms, steps of 100ms) across groups (i.e., PD-PH vs. PD-nPH). Here, the data was analyzed with a linear model, fitted for each participant independently. We assessed (1) the main effect of the delay (on the intensity of riPH) with a permutation test (5000 iterations) between slopes of the individual fit vs. zero; (2) the difference between the slopes of PD-PH vs. PD-nPH with a permutation test between the slopes of the two subgroups; (3) the main effect of group with a permutation test on the intercepts between the two subgroups.

### Study 2

#### Participants, ethics, and informed consent (study2.1)

All healthy participants had no history of neurological or psychiatric disorders. All participants provided written informed consent prior to the experiment. The study was approved by the Cantonal Ethics Committee of Geneva (Commission Cantonale d’Ethique de la Recherche sur l’Être Humain-CCER). Twenty-five healthy participants (10 women, mean age±SD: 24.6±3.7 years old; age range: 18-32 years old, Edinburg Handedness Inventory mean index: 64.8±23.7 and range: 30-100) took part in study2.1.

#### Experimental procedure (study2.1)

The experimental procedure was based on a pilot study performed in a mock scanner (Supplementary S16). Participants were blindfolded during the task and received auditory cues through earphones to start (1 beep) and to stop (2 beeps) the movement. The paradigm was implemented using an in-house software (ExpyVR, http://lnco.epfl.ch/expyvr) and Visual studio 2013 interface (Microsoft) was used to control the robotic system.

Participants underwent two runs of 12 min each, during which they repeatedly had to move the front robot for 30s with their right hand followed by 20s of rest for a total of 16 repetitions per condition (8 repetitions for the motor and touch control tasks) (Supplementary S15-S19 and Fig.S3). Synchronous and the asynchronous conditions were randomized across runs. The questionnaire was presented at the end of the scanning session and after a randomized repetition of 30s of each condition. The questionnaire was based on the pilot study (Supplementary S16-S18) and on a previous study^26^. Participants were asked to indicate on a 7-point Likert scale, how strongly they felt the sensation described by each item (from 0 = not at all, to 6 = very strong).

### Questionnaire analysis

Questionnaire data were analyzed in the same way as in study1.1. Synchrony (synchronous and asynchronous) was used as a fixed effect and the subjects as random intercepts.

#### fMRI experiment

##### fMRI data acquisition

The imaging data was acquired with a 3T Siemens Magnetom Prisma MR scanner at Campus Biotech MR Platform (Geneva). The functional data were acquired using an Echo Planar Imaging (EPI) sequence with a full brain coverage (43 continuous slices, FOV=230mm, TR=2.5s, TE=30ms, flip angle=90°, in-plane resolution=2.5×2.5mm2, slice thickness=2.5mm using a 64-channel head-coil) containing 320 volumes for the experimental runs and 160 volumes for the localizer runs. For each participant, an anatomical image was recorded using a T1-weighted MPRAGE sequence (TR=2.3s, TE=2.32 ms, Inversion time=900ms, flip angle=8°, 0.9mm isotropic voxels, 192 slices per slab and FOV=240mm).

##### fMRI data analysis

All the fMRI data analysis reported were pre-processed using SPM12 toolbox (Wellcome Departement of Cognitive Neurology, Institute of Neurology, UCL, London, UK) in Matlab (R2016b, Mathworks). Slice timing correction and spatial realignment was applied to individual functional images. The anatomical image was then co-registered with the mean functional image and segmented into grey matter, white matter and cerebro-spinal fluid (CSF) tissue. Finally, the anatomical and the functional images were normalized to the Montreal Neurological Institute (MNI) brain template. Functional images were then smoothed with a Gaussian kernel with full-width half-maximum of 6mm. Head motion was assessed based on framewise displacement (FD) calculation^69^. All participants had a mean FD value inferior to 0.50mm (mean FD=0.12±0.05 mm). The two experimental runs were filtered with a high-pass filter at 1/300 Hz to remove low frequency confounds, while the two localizers were filtered with a high-pass filter at 1/100 Hz.

##### Activation contrasts

The experimental runs and functional localizers were submitted to a general linear model (GLM) analysis. In all runs, the periods corresponding to a given robotic stimulation (i.e., synchronous, asynchronous, motor task, touch task (Supplementary S19 and Fig.S3)) and the periods corresponding to the auditory cues were modelled as separated regressors. The six realignment parameters were modelled for each run as regressors of no interest. In order to avoid confounding effects due to the amount of movement performed in each trial, the quantity of movement of the front robot (synchronous and asynchronous for the experimental runs and movement condition for the motor localizer, see above) was included as parametric modulators of each condition (see above).

Second-level analyses were performed using the first-level contrasts defined for each subject. In order to determine which brain regions were involved in sensorimotor conflicts (spatio-temporal conflict and fixed spatial conflict), the following contrasts were computed: asynchronous>motor+touch and synchronous>motor+touch. A conjunction between those two contrasts was performed to identify the regions involved in the fixed spatial sensorimotor conflicts. For the experimental runs, two sample t-tests (asynchronous>synchronous and synchronous>asynchronous) were performed to assess brain activations activated during a specific sensorimotor conflict. Results were thresholded at p<0.001 at voxel level and only the clusters surviving p<0.05 FWE-corrected for multiple comparison were reported as significant. The obtained clusters were labelled using the AAL atlas^70^ and the Anatomy toolbox^71^.

#### Lesion network mapping analysis (study2.2)

In order to identify the brain regions functionally connected to each lesion location causing PH in neurological patients, we used lesion network mapping analysis^50,72^. Briefly, this method uses normative resting state data from 151 healthy subjects obtained from the publicly available Enhanced Nathan Kline Institute Rockland Sample^51^ and uses the lesion locations as seed ROI. The fMRI acquisition parameters are described in the Supplementary S21.

##### Resting state fMRI analysis

For the pre-processing steps see above and Supplementary S21. The anatomical T1-weighted image was segmented into grey and white matter and CSF. Spatial realignment was applied to individual functional images. The six realignment parameters and their first-degree derivatives were added in addition to the averaged signals of the white matter and cerebro-spinal fluid. Subjects with the excessive motion were excluded from the analysis, this comprised 25 subjects which had a mean FD higher than 0.5mm and where more than 15% of scans were affected by movement. In total, 126 subjects were included for the analysis. Then, fMRI data was bandpass-filtered in the range of 0.008-0.09Hz.

The resting state data was analyzed using the CONN-fMRI Functional Connectivity toolbox^73^ (v.18.a, http://www.nitrc.org/projects/conn). The lesion masks were used as seed ROIs and their mean time course was extracted and correlated to all other brain voxels. Each lesion-seed yielded a brain network thresholded at p<0.001(t±3.37) with p<0.05 whole brain FWE peak level corrected. The 11 networks were then binarized and overlapped to determine the regions of shared positive and negative correlations (Fig.S5). The network overlap was thresholded at 90% (at least 10 cases out of 11) with a minimal cluster extent of 50 voxels. This procedure was repeated with the visual hallucinations (VH) lesions (Supplementary S22-S23 for further analyses).

### Study 3

#### Participants (study3.1)

Data from thirty PD patients were analyzed in this study. All patients were prospectively recruited from a sample of outpatients regularly attending to the Movement Disorders Clinic at Hospital de la Santa Creu i Sant Pau (Barcelona) based on the fulfilling of MDS new criteria for PD. Informed consent to participate in the study was obtained from all participants. The study was approved by the local Ethics Committee. Patients were diagnosed by a neurologist with expertise in movement disorders. Each patient was interviewed regarding years of formal education, disease onset, medication history, current medications, and dosage (levodopa daily dose). Motor status and stage of illness were assessed by the MDS-UPDRS-III. All participants were on stable doses of dopaminergic drugs during the 4 weeks before inclusion. Patients were included if the hallucinations remained stable during the 3 months before inclusion in the study. No participant had used or was using antipsychotic medication (Supplementary S24). Details of image acquisition and data processing are in Supplementary S25.

##### Regions of interest

The cPH-network as defined in Study 2 (right posterior middle temporal gyrus (pMTG; x =54, y=-54, z=0), the right inferior frontal gyrus (IFG; x=51, y=18, z=29) and the left ventral premotor cortex (vPMC; x=-53, y=1, z=37) was transposed bilaterally to ensure that the cPH-network is not affected by any effects of movement-related laterality of activation observed in the riPH-networks (Fig.3B). Clusters were built using FSL (https://fsl.fmrib.ox.ac.uk/fsl/). A control network was derived by shifting each region (x±0/20; y+30; z-15) of the cPH-network (Fig.S7). This approach allowed controlling for the exact same shape and number of voxels as original cPH-network areas.

##### Statistical analyses

To assess whether the functional connectivity of the cPH-network predicted if a patient was clinically classified PD-PH (or PD-nPH), we conducted a leave one out cross-validation procedure with a linear discriminant analysis (LDA) (using Caret R packages^81^). To ensure that the kappa value was above chance-level we conducted a permutation test (5000 iterations). At each iteration, functional connectivity values were permuted between sub-groups and the cross-validation procedure was repeated. Post-hoc analyses for the between group differences were performed using a permutation tests (5000 iterations) on the connection which mostly contributed to the decoding. Connectivity outliers (8.75% of all data points) were identified based on 1.5 IQR from the connectivity median value for each connection. Spearman 2-tailed correlation analyses were performed between functional connectivity within cPH-network areas and neuropsychological measures of the PD-CRS (Parkinson’s disease – Cognitive Rating Scale). Significance between the two correlations was assessed using the Steiger Tests (psych R package^76^).

All the authors declare no competing interests.

## Acknowledgments

We thank Dr. Didier Genoud and Dr. Vanessa Fleury for their contribution in recruiting patients.

## Code & Data availability

Matlab and R code, behavioral and MRI data of this study are available from the corresponding author (Olaf Blanke) upon reasonable request.

## Reference List

1. Jaspers, K. Über leibhaftige Bewusstheiten (Bewusstheitstaüschungen), ein psychopathologisches Elementarsymptom. Zeitschrift für Pathopsychologie 2, 150–161 (1913).

2. Critchley, M. The idea of a presence. Acta Psychiatrica Scandinavica 30, 155–168 (1955).

3. Arzy, S., Seeck, M., Ortigue, S., Spinelli, L. & Blanke, O. Induction of an illusory shadow person. Nature 443, 287 (2006).

4. Messner, R. The Naked Mountain. (Seattle: Cambridge University Press, 2003).

5. Geiger, J. The Third Man Factor: Surviving the Impossible. (New York: Weinstein Books, 2009).

6. Llorca, P. M. et al. Hallucinations in schizophrenia and Parkinson’s disease: an analysis of sensory modalities involved and the repercussion on patients. Scientific Reports 6, 38152 (2016).

7. Brugger, P., Regard, M. & Landis, T. Unilaterally Felt ‘Presences’: The Neuropsychiatry of One’s Invisible Doppelgänger. Neuropsychiatry, neuropsychology, and behavioral neurology 9, 114–122 (1996).

8. Arzy, S., Seeck, M., Ortigue, S., Spinelli, L. & Blanke, O. Induction of an illusory shadow person. Nature 443, 287 (2006).

9. Critchley, M. The divine banquet of the brain and other essays. (Raven Press, 1979).

10. Fénelon, G., Soulas, T., De Langavant, L. C., Trinkler, I. & Bachoud-Lévi, A.-C. Feeling of presence in Parkinson’s disease. J Neurol Neurosurg Psychiatry 82, 1219–1224 (2011).

11. Lenka, A., Pagonabarraga, J., Pal, P. K., Bejr-Kasem, H. & Kulisevsky, J. Minor hallucinations in Parkinson disease: A subtle symptom with major clinical implications. Neurology (2019) doi:10.1212/WNL.0000000000007913.

12. Ffytche, D. H. et al. The psychosis spectrum in Parkinson disease. Nat Rev Neurol 13, 81–95 (2017).

13. Fénelon, G., Soulas, T., Zenasni, F. & De Langavant, L. C. The changing face of Parkinson’s disease-associated psychosis: a cross-sectional study based on the new NINDS-NIMH criteria. Mov Disord 25, 755–759 (2010).

14. Diederich, N. J., Fénelon, G., Stebbins, G. & Goetz, C. G. Hallucinations in Parkinson disease. Nat Rev Neurol 5, 331–342 (2009).

15. Forsaa, E. B., Larsen, J. P., Wentzel-Larsen, T. & Alves, G. What predicts mortality in Parkinson disease?: a prospective population-based long-term study. Neurology 75, 1270–1276 (2010).

16. Marinus, J., Zhu, K., Marras, C., Aarsland, D. & van Hilten, J. J. Risk factors for non-motor symptoms in Parkinson’s disease. The Lancet Neurology 17, 559–568 (2018).

17. Goetz, C. G., Fan, W., Leurgans, S., Bernard, B. & Stebbins, G. T. The malignant course of ‘benign hallucinations’ in Parkinson disease. Arch. Neurol. 63, 713–716 (2006).

18. Pagonabarraga, J. et al. Minor hallucinations occur in drug-naive Parkinson’s disease patients, even from the premotor phase. Mov. Disord. 31, 45–52 (2016).

19. Kulick, C. V., Montgomery, K. M. & Nirenberg, M. J. Comprehensive identification of delusions and olfactory, tactile, gustatory, and minor hallucinations in Parkinson’s disease psychosis. Parkinsonism Relat. Disord. 54, 40–45 (2018).

20. Fernandez, H. H. et al. Scales to assess psychosis in Parkinson’s disease: Critique and recommendations. Movement Disorders 23, 484–500 (2008).

21. Ravina, B. et al. Diagnostic criteria for psychosis in Parkinson’s disease: Report of an NINDS, NIMH Work Group. Movement Disorders 22, 1061–1068 (2007).

22. Holroyd, S., Currie, L. & Wooten, G. F. Prospective study of hallucinations and delusions in Parkinson’s disease. Journal of Neurology, Neurosurgery & Psychiatry 70, 734–738 (2001).

23. Weil, R. S. et al. Visual dysfunction in Parkinson’s disease. Brain 139, 2827–2843 (2016).

24. Bejr-Kasem, H. et al. Disruption of the default mode network and its intrinsic functional connectivity underlies minor hallucinations in Parkinson’s disease. Mov. Disord. 34, 78–86 (2019).

25. H Hecaen & De Ajuriaguerra. Misconstructions and hallucinations with respect to the body image; integration and disintegration of somatognosi. L’ Evolution Psychiatrique 745–750 (1952).

26. Blanke, O. et al. Report Neurological and Robot-Controlled Induction of an Apparition. Current Biology 24, 2681–2686 (2014).

27. Weiskrantz, L., Elliott, J. & Darlington, C. Preliminary observations on tickling oneself. Nature 230, 598–599 (1971).

28. Blakemore, S. J., Wolpert, D. & Frith, C. Why can’t you tickle yourself? Neuroreport 11, R11–16 (2000).

29. Ehrsson, H. H., Holmes, N. P. & Passingham, R. E. Touching a rubber hand: feeling of body ownership is associated with activity in multisensory brain areas. J Neurosci 25, 10564–10573 (2005).

30. Pozeg, P., Rognini, G., Salomon, R. & Blanke, O. Crossing the Hands Increases Illusory Self-Touch. PLoS One 9, (2014).

31. Shergill, S. S., Samson, G., Bays, P. M., Frith, C. D. & Wolpert, D.M. Evidence for sensory prediction deficits in schizophrenia. Am J Psychiatry 162, 2384–2386 (2005).

32. Blakemore, S.-J., Wolpert, D. M. & Frith, C.D. Central cancellation of self-produced tickle sensation. Nat Neurosci 1, 635–640 (1998).

33. Blakemore, S.-J., Wolpert, D. M. & Frith, C.D. Abnormalities in the awareness of action. Trends in Cognitive Sciences 6, 237–242 (2002).

34. Wolpert, D. M., Ghahramani, Z. & Jordan, M.I. An internal model for sensorimotor integration. Science 269, 1880–1882 (1995).

35. Miall, R. C. & Wolpert, D.M. Forward Models for Physiological Motor Control. Neural Networks 9, 1265–1279 (1996).

36. Corlett, P. R. et al. Disrupted prediction-error signal in psychosis: evidence for an associative account of delusions. Brain 130, 2387–2400 (2007).

37. Fletcher, P. C. & Frith, C.D. Perceiving is believing: a Bayesian approach to explaining the positive symptoms of schizophrenia. Nat. Rev. Neurosci. 10, 48–58 (2009).

38. Ford, J. M. & Mathalon, D.H. Electrophysiological evidence of corollary discharge dysfunction in schizophrenia during talking and thinking. Journal of Psychiatric Research 38, 37–46 (2004).

39. Conte, A., Khan, N., Defazio, G., Rothwell, J. C. & Berardelli, A. Pathophysiology of somatosensory abnormalities in Parkinson disease. Nature Reviews Neurology 9, 687–697 (2013).

40. Pagonabarraga, J. et al. Neural correlates of minor hallucinations in non-demented patients with Parkinson’s disease. Parkinsonism & Related Disorders 20, 290–296 (2014).

41. Hara, M. et al. A novel manipulation method of human body ownership using an fMRI-compatible master-slave system. Journal of Neuroscience Methods 235, 25–34 (2014).

42. Leube, D. T. et al. The neural correlates of perceiving one’s own movements. NeuroImage 20, 2084–2090 (2003).

43. Sperduti, M., Delaveau, P., Fossati, P. & Nadel, J. Different brain structures related to self- and external-agency attribution: a brief review and meta-analysis. Brain Struct Funct 216, 151–157 (2011).

44. David, N., Newen, A. & Vogeley, K. The ‘’ sense of agency “and its underlying cognitive and neural mechanisms. Consciousness and Cognition 17, 523–534 (2008).

45. Blakemore, S. J., Wolpert, D. M. & Frith, C. D. Central cancellation of self-produced tickle sensation. Nature neuroscience 1, 635–640 (1998).

46. Yomogida, Y. et al. The neural basis of agency: an fMRI study. Neuroimage 50, 198–207 (2010).

47. Ehrsson, H. H., Holmes, N. P. & Passingham, R. E. Touching a rubber hand: feeling of body ownership is associated with activity in multisensory brain areas. The Journal of neuroscience □ : the official journal of the Society for Neuroscience 25, 10564–73 (2005).

48. Farrer, C. et al. Modulating the experience of agency: A positron emission tomography study. NeuroImage 18, 324–333 (2003).

49. Blakemore, S. J. & Sirigu, A. Action prediction in the cerebellum and in the parietal lobe. Experimental Brain Research 153, 239–245 (2003).

50. Boes, A. D. et al. Network localization of neurological symptoms from focal brain lesions. Brain 138, 3061–3075 (2015).

51. Nooner, K. B. et al. The NKI-Rockland Sample: A Model for Accelerating the Pace of Discovery Science in Psychiatry. Front Neurosci 6, (2012).

52. Friston, K. J. The disconnection hypothesis. Schizophr. Res. 30, 115–125 (1998).

53. Friston, K., Brown, H. R., Siemerkus, J. & Stephan, K. E. The dysconnection hypothesis (2016). Schizophrenia Research 176, 83–94 (2016).

54. Frith, C. The neural basis of hallucinations and delusions. C. R. Biol. 328, 169–175 (2005).

55. Aarsland, D., Hutchinson, M. & Larsen, J. P. Cognitive, psychiatric and motor response to galantamine in Parkinson’s disease with dementia. Int J Geriatr Psychiatry 18, 937–941 (2003).

56. Ramirez-Ruiz, B., Junque, C., Marti, M.-J., Valldeoriola, F. & Tolosa, E. Cognitive changes in Parkinson’s disease patients with visual hallucinations. Dement Geriatr Cogn Disord 23, 281–288 (2007).

57. Morgante, L. et al. Psychosis associated to Parkinson’s disease in the early stages: relevance of cognitive decline and depression. J. Neurol. Neurosurg. Psychiatry 83, 76–82 (2012).

58. Pagonabarraga, J. et al. Parkinson’s disease-cognitive rating scale: a new cognitive scale specific for Parkinson’s disease. Mov. Disord. 23, 998–1005 (2008).

59. Blakemore, S. J. & Frith, C. Self-awareness and action. Curr. Opin. Neurobiol. 13, 219–224 (2003).

60. Friston, K. J. Theoretical neurobiology and schizophrenia. Br. Med. Bull. 52, 644–655 (1996).

61. Goetz, C. G., Vaughan, C. L., Goldman, J. G. & Stebbins, G. T. I finally see what you see: Parkinson’s disease visual hallucinations captured with functional neuroimaging. Mov. Disord. 29, 115–117 (2014).

62. Meppelink, A. M. et al. Impaired visual processing preceding image recognition in Parkinson’s disease patients with visual hallucinations. Brain 132, 2980–2993 (2009).

63. Weintraub, D., Kales, H. C. & Marras, C. The Danger of Not Treating Parkinson Disease Psychosis-Reply. JAMA Neurol 73, 1156–1157 (2016).

64. Rana, A. Q., Vaid, H. M., Edun, A., Dogu, O. & Rana, M. A. Relationship of dementia and visual hallucinations in tremor and non-tremor dominant Parkinson’s disease. J. Neurol. Sci. 323, 158–161 (2012).

65. Reijnders, J. S. a. M., Ehrt, U., Lousberg, R., Aarsland, D. & Leentjens, A. F.G. The association between motor subtypes and psychopathology in Parkinson’s disease. Parkinsonism Relat. Disord. 15, 379–382 (2009).

66. Bates, D., Mächler, M., Bolker, B. & Walker, S. Fitting Linear Mixed-Effects Models Using lme4. Journal of Statistical Software 67, 1–48 (2015).

67. Kuznetsova, A., Brockhoff, P. B. & Christensen, R. H. B. lmerTest Package: Tests in Linear Mixed Effects Models. Journal of Statistical Software 82, 1–26 (2017).

68. Luo, D, Ganesh, S & Koolaard, J. predictmeans: Calculate Predicted Means for Linear Models, http://cran.r-project.org/package=predictmeans. (2014).

69. Power, J. D., Barnes, K. A., Snyder, A. Z., Schlaggar, B. L. & Petersen, S. E. Spurious but systematic correlations in functional connectivity MRI networks arise from subject motion. Neuroimage 59, 2142–2154 (2012).

70. Tzourio-Mazoyer, N. et al. Automated Anatomical Labeling of Activations in SPM Using a Macroscopic Anatomical Parcellation of the MNI MRI Single-Subject Brain. NeuroImage 15, 273–289 (2002).

71. Eickhoff, S. B. et al. A new SPM toolbox for combining probabilistic cytoarchitectonic maps and functional imaging data. NeuroImage 25, 1325–1335 (2005).

72. Fox, M. D. Mapping Symptoms to Brain Networks with the Human Connectome. New England Journal of Medicine 379, 2237–2245 (2018).

73. Whitfield-Gabrieli, S. & Nieto-Castanon, A. Conn: a functional connectivity toolbox for correlated and anticorrelated brain networks. Brain Connect 2, 125–141 (2012).

74. Kuhn, M. caret: Classification and Regression Training. Astrophysics Source Code Library ascl:1505.003 (2015).

75. Friedman, J., Hastie, T. & Tibshirani, R. Regularization Paths for Generalized Linear Models via Coordinate Descent. J Stat Softw 33, 1–22 (2010).

76. Revelle, W. R. psych: Procedures for Personality and Psychological Research. (2017).

