## Supplemental information for "Sensorimotor hallucinations in Parkinson’s disease"

**Affiliations**

#### *Authors' contributions*

FB designed Study 1 and 3, collected & analyzed data, conducted clinical interviews, wrote paper; EB designed Study 2, collected & analyzed data, wrote paper; J. Potheegadoo collected data, designed questionnaire for semi-structured interview, conducted clinical interviews and clinical evaluations for study 1; GS analyzed data for study 3; J. Pagonabarraga, HB and JK recruited patients, conducted clinical interviews, collected data for study 3; MA and NF analyzed data for study 2; MB collected data, conducted clinical interviews and clinical evaluations for study 1; MF collected data for study 1; SK coordinated the recruitment for study 1; SMH designed and conducted clinical interviews for study 3; FS collected data for study 3; MH designed and developed the robotic systems; JH, JG, PB recruited patients and conducted clinical evaluations for study 1; DV designed study 2; PK designed study 1; GR and OB designed study 1, 2 and 3, wrote paper. All authors provided critical revisions and approved the final version of the paper for submission.

All the authors declare no competing interests.

#### *Acknowledgments*

We thank Dr. Didier Genoud and Dr. Vanessa Fleury for their contribution in recruiting patients.

*** These authors equally contributed to the work**

**Co-corresponding authors**

Olaf Blanke
Bertarelli Chair in Cognitive Neuroprosthetics
Center for Neuroprosthetics & Brain Mind Institute
School of Life Sciences
Campus Biotech
Swiss Federal Institute of Technology
Ecole Polytechnique Fédérale de Lausanne (EPFL)
CH – 1012 Geneva


Jaime Kulisevsky
Movement Disorders Unit
Neurology Department
Hospital de la Santa Creu i Sant Pau
Mas Casanovas 90, 08041
Barcelona, Spain

Funding: Carigest SA, Swiss National Science Foundation (3100A0-112493), Parkinson Suisse, Bertarelli Foundation to Olaf Blanke, CIBERNED (Carlos III Institute) to Jaime Kulisevsky, JSPS Fund for the Promotion of Joint International Research (Fostering Joint International Research) (17KK0003) to Masayuki Hara.

Keywords: Parkinson's disease, Hallucinations, Sensorimotor, fMRI, cognitive decline

### Study 1

### Study 1.1: Robot-induced presence hallucinations (riPH) in patients with PD

#### *Supplementary S1*

**Participants: Inclusion/Exclusion criteria**

Participants in the present study consisted of patients with PD and the symptom of PH (PD-PH, n=13), patients with PD without the symptom of PH (PD-nPH, n=13), and age-matched healthy controls (HC, n=21). Demographic and clinical data are summarized in Table S1. Patients with cognitive impairments (defined as a MoCA score[1] lower then 24[2]), treated with neuroleptics, affected by other central neurological co-morbidities, affected by psychiatric co-morbidities unrelated to PD, and patients with recent (< one month) changes in their medical treatment were not included in the study. The HC included in the study never experienced PH, did not suffer from a neurological or psychiatric disease, and had no objective sign of cognitive impairment.

#### *Supplementary S2*

**Demographic and disease-related variables**

For every PD patient, the doses of anti-parkinsonian medication were converted to the levodopa equivalent daily dose[3]. The severity of motor symptoms was assessed by the score at the Movement Disorders Society - Unified Parkinson Disease Rating Scale (MDS-UPDRS) - part III (Goetz et al., 2008), in "ON" state. In addition, impulsive-compulsive disorders were assessed by the score at the "Questionnaire for Impulsive-Compulsive Disorders in Parkinson's Disease" (QUIP-RS[4]). We also assessed for PD-PH, PD-nPH, and HC apathy scale[5] and the risk for psychosis

(Prodromal Questionnaire PQ-16[6]; which was divided in part I (hallucinations, and negative symptoms-like experiences), and part II (level of distress linked to the experiences). Hallucinations were assessed with a semi-structured interview adapted from the psychosensory hallucinations Scale (PSAS) for Schizophrenia and Parkinson's disease[7]. Next to PH, we also inquired about other hallucinations possibly experienced by patients with PD, e.g. passage hallucinations (i.e., animal, person or indefinite object passing in the peripheral visual field), visual illusion and complex hallucinations (structured visual, auditory or tactile hallucinations) as well as delusional ideas.

| | **PD-PH (N = 13)** | **PD-nPH (N = 13)** | ***p-values*** |
|---|---|---|---|
| Age | 60.69 ± 13.19 | 65.69 ± 7.60 | 0.25 |
| Gender | 9 (M) | 4 (M) | 0.05 ($\chi2$) |
| UPDRS-III | 20 ± 12.09 | 19 ± 17.51 | 0.87 |
| MoCA | 26.85 ± 1.82 | 28.15 ± 1.57 | 0.08 |
| PQ16 | 4.00 ± 2.00 | 0.69 ± 1.32 | < 0.001 |
| PQ16-2 | 3.54 ± 4.86 | 1.08 ± 2.63 | 0.1 |
| Apathy | 12.69 ± 8.06 | 10.23 ± 4.64 | 0.37 |
| LEDD (mg/day) | 727.77 ± 410.46 | 786.23 ± 657.23 | 0.8 |
| Disease Duration (years) | 9.46 ± 4.22 | 9.38 ± 5.72 | 0.5 |

**Table S1. Clinical variables between PD-PH and PD-nPH.**

| | **PD-PH (N = 13)** | **HC (N = 21)** | ***p-values*** |
|---|---|---|---|
| Age | 60.69 ± 13.19 | 66.90 ± 5.75 | 0.06 |
| Gender | 9 (M) | 11 (M) | 0.9 ($\chi2$) |
| MoCA | 26.85 ± 1.82 | 28.52 ± 1.03 | <0.001 |
| PQ16 | 4.00 ± 2.00 | 0.24 ± 0.44 | <0.001 |
| PQ16-2 | 3.54 ± 4.86 | 0 ± 0 | <0.001 |
| Apathy | 12.69 ± 8.06 | 6.33 ± 4.05 | 0.01 |

**Table S2. Clinical variables between PD-PH and HC.**

### *Supplementary S3*

**Experimental procedure**

Each PD patient underwent study1 at a similar time (10am), after having received their usual anti-parkinsonian medication and were in their "best ON" state for the whole duration of study1 as well as the psychological and neuropsychological assessments[8]. To investigate the riPH in patients with PD (and HC), we used the same experimental setup and device as our previous research[9]. The robotic stimulation was administered through a robotic system[10] that has previously been used to induce the PH and other bodily illusions in healthy subjects[9]. The experimental design consisted in factors Synchrony (synchronous/asynchronous), Side (most/less affected) and Group (PD-PH/PD-nPH).

### *Supplementary S4*

***Questionnaire results: PH***

Detailed ratings for all questions can be seen on Table S3 (below).

**riPH** *("I felt as if someone was close-by")*

*PD-PH vs. PD-nPH.* No main effect of Side (permutation p-value=0.37). No interactions were observed, all permutation p-values>0.05.

*PD-PH vs. HC.* By comparing PD-PH and HC, we confirmed the importance of conflicting sensorimotor stimulation to induced PH, as both groups gave higher PH ratings in the asynchronous versus synchronous condition (p-value=0.033). The intensity of riPH ratings did not differ statistically between PD-PH and HC (permutation p-value=0.48). The Side did not significantly modulate the riPH ratings (permutation p-value=0.38). No interactions were observed, all permutation p-values>0.05.

### *Supplementary S5*

***Questionnaire results: Other robot-induced perceptions***

*Passivity experience ("I felt as if someone else was touching my back.").*

*PD-PH vs. PD-nPH.* The two sub-groups of patients did not report difference in passivity experiences in the asynchronous condition (permutation p-value = 0.1, main effect of Synchrony), the ratings did not differ significantly between the groups of patients (permutation p-value = 0.38, main effect of Group), and the Side did not modulate the passivity experience (permutation p-value=0.41). No interactions were observed, all permutation p-values>0.05.

*PD-PH vs. HC.* We observed a trend for asynchronous condition to induce higher passivity experiences in the asynchronous condition (permutation p-value=0.06, main effect of Synchrony). The ratings were not statistically different between the groups (permutation p-value=0.86, main effect of Group). The Side modulated the passivity experience (permutation p-value<0.01). No interactions were observed, all other permutation p-values>0.05.

*Self-touch ("I felt as if I was touching my back.").*

*PD-PH vs. PD-nPH.* In line with previous work[9], the two sub-groups of patients reported higher self-touch experiences in the synchronous condition (permutation p-value=0.043, main effect of Synchrony). The ratings did not differ significantly neither between the groups of patients (permutation p-value=0.65, main effect of Group), nor between the Side (permutation p-value=0.51). No interactions were observed, all other permutation p-values>0.05.

*PD-PH vs. HC.* We observed that participants reported a trend for higher self-touch experiences in the synchronous condition (permutation p-value=0.054, main effect of Synchrony). The rating did not differ significantly neither between the groups (permutation p-value=0.92, main effect of Group), nor between the Side (permutation p-value=0.4). No interactions were observed (all other permutation p-values>0.05).

*Loss of agency ("I felt as if I was not controlling my movements or actions.").*

*PD-PH vs. PD-nPH.* The robotic stimulation was associated with a stronger loss of agency in PD-PH than PD-nPH (permutation p-value=0.045, main effect of Group). Neither the sensorimotor conditions (permutation p-value=0.26, main effect of

Synchrony), nor the Side (permutation p-value=0.67) modulated significantly the rating. No interactions were observed, all other permutation p-values>0.05.

*PD-PH vs. HC.* No statistical difference was observed between the two sub-groups (permutation p-value=0.073, main effect of Group). Neither the sensorimotor conditions (permutation p-value=0.6, main effect of Synchrony) nor the Side (permutation p-value=0.28) modulated significantly the ratings. No interactions were observed, all other permutation p-values>0.05.

*Bodily sensations* (*"I felt as if I had two bodies") (Control item 1*).

*PD-PH vs. PD-nPH.* The robotic stimulation did not modulate significantly this bodily sensation (permutation p-value=0.98, main effect of Synchrony), we did not observe statistically significant differences between the two sub-groups (permutation p-value=0.26, main effect of Group), and did not observed a difference due to the Side (permutation p-value=0.88). No interactions were observed, all other permutation p-values>0.05.

*PD-PH vs. HC.* The robotic stimulation neither modulated this bodily sensation (permutation p-value=0.85, main effect of Synchrony) nor did the two sub-groups (permutation p-value=0.79, main effect of Group), and we did not observe a difference due to the Side (permutation p-value=0.71). No interactions were observed, all other permutation p-values>0.05.

*Control Question ("I felt someone was standing in front of me.") (Control item 2).*

For each of the three sub-groups, all the raw ratings were zeros for the question front-PH (permutation p-value=1).

| Question | Group | Synchrony | Side | Mean | Standard Deviation |
|---|---|---|---|---|---|
| PH | PD-PH | Async | Less affected side | 1.67 | 2.31 |
| PH | PD-PH | Async | Most affected side | 2.23 | 2.17 |
| PH | PD-PH | Sync | Less affected side | 0.42 | 0.9 |
| PH | PD-PH | Sync | Most affected side | 1.31 | 1.93 |
| PH | PD-nPH | Async | Less affected side | 0.62 | 1.56 |
| PH | PD-nPH | Async | Most affected side | 0.23 | 0.83 |
| PH | PD-nPH | Sync | Less affected side | 0 | 0 |
| PH | PD-nPH | Sync | Most affected side | 0 | 0 |
| PH | HC | Async | Less affected side | 1.05 | 1.96 |
| PH | HC | Async | Most affected side | 1.33 | 2.15 |
| PH | HC | Sync | Less affected side | 0.52 | 1.66 |
| PH | HC | Sync | Most affected side | 0.86 | 1.93 |

| Question | Group | Synchrony | Side | Mean | Standard Deviation |
|---|---|---|---|---|---|
| Loss of agency | PD-PH | Async | Less affected side | 1.83 | 2.25 |
| Loss of agency | PD-PH | Async | Most affected side | 2.25 | 2.14 |
| Loss of agency | PD-PH | Sync | Less affected side | 1.17 | 1.7 |
| Loss of agency | PD-PH | Sync | Most affected side | 1.75 | 1.71 |
| Loss of agency | PD-nPH | Async | Less affected side | 0.85 | 1.63 |
| Loss of agency | PD-nPH | Async | Most affected side | 0.54 | 0.97 |
| Loss of agency | PD-nPH | Sync | Less affected side | 0.08 | 0.28 |
| Loss of agency | PD-nPH | Sync | Most affected side | 0.23 | 0.6 |
| Loss of agency | HC | Async | Less affected side | 0.48 | 1.25 |
| Loss of agency | HC | Async | Most affected side | 0.81 | 1.47 |
| Loss of agency | HC | Sync | Less affected side | 0.24 | 0.89 |
| Loss of agency | HC | Sync | Most affected side | 0.9 | 1.7 |

| Question | Group | Synchrony | Side | Mean | Standard Deviation |
|---|---|---|---|---|---|
| Passivity experience | PD-PH | Async | Less affected side | 2.33 | 2.31 |
| Passivity experience | PD-PH | Async | Most affected side | 3.08 | 2.43 |
| Passivity experience | PD-PH | Sync | Less affected side | 1.25 | 2.05 |
| Passivity experience | PD-PH | Sync | Most affected side | 2.08 | 2.14 |
| Passivity experience | PD-nPH | Async | Less affected side | 2.54 | 2.37 |
| Passivity experience | PD-nPH | Async | Most affected side | 1.77 | 2.05 |
| Passivity experience | PD-nPH | Sync | Less affected side | 1.54 | 2.22 |
| Passivity experience | PD-nPH | Sync | Most affected side | 1.38 | 1.94 |
| Passivity experience | HC | Async | Less affected side | 1.81 | 2.5 |
| Passivity experience | HC | Async | Most affected side | 3.29 | 2.31 |
| Passivity experience | HC | Sync | Less affected side | 1.29 | 2.22 |
| Passivity experience | HC | Sync | Most affected side | 2.33 | 2.61 |

| Question | Group | Synchrony | Side | Mean | Standard Deviation |
|---|---|---|---|---|---|
| Self-touch | PD-PH | Async | Less affected side | 1.92 | 2.35 |
| Self-touch | PD-PH | Async | Most affected side | 1.38 | 1.66 |
| Self-touch | PD-PH | Sync | Less affected side | 2.08 | 2.27 |
| Self-touch | PD-PH | Sync | Most affected side | 3 | 2.24 |
| Self-touch | PD-nPH | Async | Less affected side | 0.85 | 1.57 |
| Self-touch | PD-nPH | Async | Most affected side | 0.85 | 1.57 |
| Self-touch | PD-nPH | Sync | Less affected side | 1.46 | 2.37 |
| Self-touch | PD-nPH | Sync | Most affected side | 1.92 | 2.56 |
| Self-touch | HC | Async | Less affected side | 2.38 | 2.69 |
| Self-touch | HC | Async | Most affected side | 1.86 | 2.46 |
| Self-touch | HC | Sync | Less affected side | 2.43 | 2.84 |
| Self-touch | HC | Sync | Most affected side | 2.81 | 2.84 |

| Question | Group | Synchrony | Side | Mean | Standard Deviation |
|---|---|---|---|---|---|
| PH front | PD-PH | Async | Less affected side | 0 | 0 |
| PH front | PD-PH | Async | Most affected side | 0 | 0 |
| PH front | PD-PH | Sync | Less affected side | 0 | 0 |
| PH front | PD-PH | Sync | Most affected side | 0 | 0 |
| PH front | PD-nPH | Async | Less affected side | 0 | 0 |
| PH front | PD-nPH | Async | Most affected side | 0 | 0 |
| PH front | PD-nPH | Sync | Less affected side | 0 | 0 |
| PH front | PD-nPH | Sync | Most affected side | 0 | 0 |
| PH front | HC | Async | Less affected side | 0 | 0 |
| PH front | HC | Async | Most affected side | 0 | 0 |
| PH front | HC | Sync | Less affected side | 0 | 0 |
| PH front | HC | Sync | Most affected side | 0 | 0 |

| Question | Group | Synchrony | Side | Mean | Standard Deviation |
|---|---|---|---|---|---|
| Control | PD-PH | Async | Less affected side | 0.25 | 0.62 |
| Control | PD-PH | Async | Most affected side | 0.54 | 1.2 |
| Control | PD-PH | Sync | Less affected side | 0.25 | 0.87 |
| Control | PD-PH | Sync | Most affected side | 0.38 | 0.96 |
| Control | PD-nPH | Async | Less affected side | 0 | 0 |
| Control | PD-nPH | Async | Most affected side | 0 | 0 |
| Control | PD-nPH | Sync | Less affected side | 0 | 0 |
| Control | PD-nPH | Sync | Most affected side | 0 | 0 |
| Control | HC | Async | Less affected side | 0.19 | 0.87 |
| Control | HC | Async | Most affected side | 0.19 | 0.87 |
| Control | HC | Sync | Less affected side | 0 | 0 |
| Control | HC | Sync | Most affected side | 0.24 | 1.09 |

**Table S3. Mean ratings for all questions, and experimental conditions**

### Supplementary S6

***Post-experiment debriefing: riPH mimic sPH (in PD-PH)***

*Patients reports.* One PD-PH patient reported that he could feel the robot-induced presence on the side (not on the back) and added (after being asked to compare s- and riPH) "it is slightly similar, but it is not exactly the same because the presence (symptomatic) is all of a sudden, while here (the riPH) it is built-up". Although, the riPH was strong felt, another PD-PH patient noted that the PH lacked some aspects of his symptomatic PH (sPH). He described that "when I feel the symptomatic PH it's like a chewing gum with a lot of taste, while here (the riPH) it was still like chewing gum but without the taste". Another PD-PH patient compared his riPH to "an adrenaline rush. Like something or someone was behind me, although there is not the possibility to have someone behind" and "I really had the impression that someone was doing something behind me". Another PD-PH patient reported that "I honestly have the impression to have someone behind me". Just after the stimulation and removal of the blindfold she added "I was surprised to see you all in front of me".

### Supplementary S7

***Post-experiment debriefing: Spatial location of the riPH***

We further determined the experienced spatial location of the riPH and whether this differed across the three participant groups. Analyzing all trials for which a participant positively rated the PH during the robotic procedure (i.e., value > 0 on Likert scale) we found that PD-PH patients reported a higher number of lateralized riPH (n=24; i.e. instances of a riPH with a value > 0; across all trials and conditions) then HC (n=18) (Chi-square: p-value = 0.003, $\chi^2(1)=9$) and PD-nPH (n=3) (Chi-square: p-value = 0.001, $\chi^2(1)=11.26$), Table S2). PD-PH reported riPH either to the side (n=14) or behind them (n=6), with no predominant location (Chi-square: p-value = 0.11, $\chi^2(1)=3.22$), while HC predominantly reported riPH behind them (n=14, and n=2 lateralized) (Chi-square: p-value = 0.006, $\chi^2(1)=9$). The most affected side did not influence the location of the riPH (all p-values>0.05). The very few instances in PD-nPH patients did not differ (behind: n=2; lateralized: n=1) (Chi-square: p-value = 1, $\chi^2(1)=0.33$). These data show that similarly to the sPH, PD-PH patients experienced riPH more often on the side, even if the tactile feedback was provided on the back, differing from HC, who always reported the location of the robot-induced presence behind them. Debriefing data also suggest that 38% of PD-PH patients report robotic-induced PH that are associated with a state that is comparable in intensity to sPH. Interestingly, these robotic-induced sPH only occurred in the asynchronous stimulation condition.

### *Supplementary S8*

***sPH in PD-PH (semi-structured interview data).***

Previous studies observed that most patients with PD who experience PH report them as neutral, as not distressing (except when it occurred for the first time), and usually short-lasting. Moreover, PH are typically felt beside or behind the patient's body (rarely also in an adjacent room)[11]. In the current study, the semi-structured interview data confirmed that sPH in PD-PH patients were in 54% neutral or positive and were in 62% of undetermined gender. In 69% the presence was either felt on the side of the patient's body and/or on the back (for other variables see Table S2). Collectively, these results are compatible with previously reported sPH in PD. Overall sPH were not predominantly located in one spatial position. That is, ~38% of the patients experienced the sPH on either sides (not simultaneously) and/or in the back, confirming that sPH are not associated with the predominantly affected side of the disease [11] (Table S4 for details).

| | Number of patients | % |
|---|---|---|
| **PH Valence** | | |
| • Positive/Neutral | 7 | 54 |
| • Negative | 6 | 46 |
| **PH Gender** | | |
| • Female only | 2 | 15 |
| • Male only | 1 | 8 |
| • Both sex | 2 | 15 |
| • Undetermined | 8 | 62 |
| **PH Lateralisation** | | |
| • Side only | 6 | 46 |
| • Back only | 3 | 23 |
| • Back and Side | 2 | 15 |
| • Front | 1 | 8 |
| • Other room | 5 | 38 |
| **Occurrence (moment)** | | |
| • Day | 3 | 23 |
| • Night | 4 | 31 |
| • Anytime | 6 | 46 |
| **Occurrence (place)** | | |
| • Home only | 8 | 62 |
| • Outside home only | 0 | 0 |
| • Both | 6 | 46 |
| **Distance of PH** | | |
| • Less than 1m | 5 | 38 |
| • More than 1m | 8 | 62 |

**Table S4. Phenomenology of the symptomatic PH in PD.**

### Study 1.2: riPH in PD-PH patients depend on sensorimotor delay

#### *Supplementary S9*

#### *Participants*

The same participants of study1.1 took part in study1.2. In total, 10 PD-PH and 12 PD-nPH and 21 HC participated to study1.2. Because of fatigue and/or tremor, two PD-PH and one PD-nPH could not participate in study1.2. One PD-PH and one PD-nPH were excluded from the analysis because they performed less than 18 trials (i.e. one session) before definitively interrupting the experiment, due to fatigue and/or excessive tremor.

#### *Supplementary S10*

#### *Experimental procedure*

For each patient, the task of study1.2 was done exclusively with the hand that was most affected by PD. HC did the task with their dominant hand. Each participant was asked to perform three sessions; each session consisted of 18 trials (3 repetitions per delay). In total, each delay was repeated 9 times. The overall experiment lasted approximately 20 minutes. Between each session, the participant could take a break according to his/her needs. One PD-PH patient performed longer sessions. In total, PD-PH completed 57.8 ± 16.9 (mean ± SD) trials, PD-nPH completed 45 ± 12.8 (mean ± SD) trials, and HC completed 53.3 ± 3.91 (mean ± SD) trials. No statistically difference across groups was observed (Welch two Sample t-test, two-tailed): PD-PH vs. PD-nPH: $t(17) = 1.97$, p-value = 0.065; PD-PH vs. HC: $t(-9) = 0.89$, p-value = 0.39.

#### *Supplementary S11*

#### *Degree of sensorimotor conflict modulates riPH*

Study1.2 confirmed that PD-PH patients experienced stronger riPH than PD-nPH patients (main effect of group: permutation p-value = 0.016; Fig.1D). Comparing the intensity of riPH between PD-PH and HC, we observed that PD-PH patients have a stronger bias in experiencing riPH than HC (main effect of group: permutation p-value=0.046t), and that the intensity of riPH increased with increasing delays for both groups (main effect of delay: p-value<0.001, two-tailed permutation test; Fig.

S1). Although PD-PH patients have a stronger bias in riPH, there was no significant difference in sensitivity (slope) between these two groups (p-value=0.6, two-tailed permutation test). On average PD-PH rated the riPH for the delays: 0ms: 35.1 ± 33.8 (percentage mean ± SD), 100ms: 34.8 ± 33.8, 200ms: 45.7 ± 38, 300ms: 39.8 ± 37, 400ms: 48.1 ± 39.5, 500ms: 48.4 ± 43.3. PD-nPH rated on average the riPH: 0ms: 6.75 ± 16.1, 100ms: 5.56 ± 19.2, 200ms: 8.33 ± 25.6, 300ms: 7.41 ± 22.4, 400ms: 8.33 ± 28.9, 500ms: 7.41 ± 25.7. HC rated on average the ri PH: 0ms: 14.3 ± 26.3, 100ms: 14.8 ± 28, 200ms: 15.3 ± 29.1, 300ms: 15.2 ± 28.3, 400ms: 23.8 ± 38.9, 500ms: 23.5 ± 34.9.

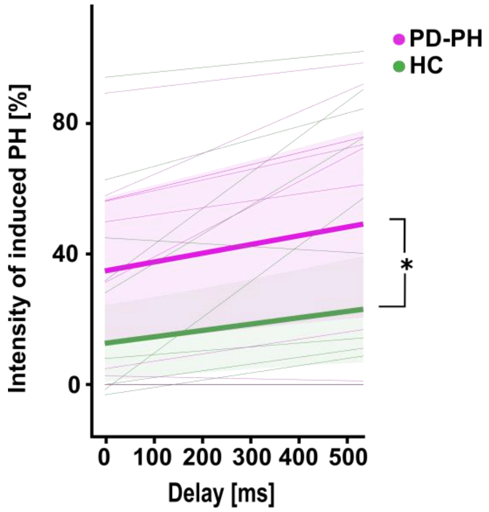


**Figure S1. riPH (PD patient & HC). A.** Study1.2. riPH were modulated by delay (permutation p-value<0.001) and PD-PH vs. HC had a stronger bias in experiencing riPH. The thicker lines indicates the mean of the fitted models, the thinner lines indicate the individual fit, and the shaded are indicates the 95% confidence interval.

### *Supplementary S12*

#### *Movement analysis*

To assess whether the spatio-temporal pattern of the movement could explain the difference in rating of the riPH among groups, we calculated: i) the Inter-poke-interval (time between the end of the touch on the back of poke *n* and the beginning of the following touch – poke *n+1*), ii) duration of the poke and iii) the spatial distance between poke *n* and poke *n+1*. Data were analyzed with linear mixed effects models lme4 and lmerTest both R packages[12,13]. The significance of fixed effects was estimated with a permutation test (5000 iterations; predicted mean R package).

*Inter-poke-interval (ipi).* To assess the temporal aspects of the sensorimotor integration we computed the ipi for each individual and for each trial independently. Models were performed on the ipi for each subject, with Groups (i.e., PD-PH vs. PD-nPH; PD-PH vs. HC as fixed effects, and random intercepts for the participant.

*Poke duration.* To assess a second temporal aspects of the sensorimotor integration we computed the duration of each poke, for each individual and for each trial independently. Models were performed on the duration for each subject, with Groups (i.e., PD-PH vs. PD-nPH; PD-PH vs. HC) as fixed effects, and random intercepts for the participant.

*Spatial distance between pokes.* To further investigate the spatial aspects of the sensorimotor integration we computed the Euclidean distance between pokes for each trial and subject. Models were performed on each distance values for each subject, with Groups (i.e., PD-PH vs. PD-nPH and PD-PH vs. HC) as fixed effects, and random intercepts for the participant.

### ***Supplementary S13***

#### ***Movement analysis***

Are bias and delay effect related to differences in the upper arm movements of PD-PH vs. PD-nPH patients during the robotic procedure? During Study1.2 we measured the movements performed by all participants, allowing us to analyze whether PD-PH, PD-nPH, and HC moved differently, calculating the inter-poke-interval (i.e., time between the end of the touch on the back (poke *n*) and the beginning of the following poke *n+1*) and the spatial distance between successive pokes (poke *n* and poke *n+1*).

The analysis of the movement data of Study1.2 exclude differences in movement patterns (neither temporal nor spatial aspects) between the two sub-groups of patients. No difference in the inter-poke-interval between PD-PH and PD-nPH (permutation p-value = 0.29). Average duration of the inter-poke-interval for PD-PH was 2.06±1.97 seconds (mean ± SD) and 1.55±2.26 sec for PD-nPH (mean ± SD). The duration of each poke did not different between PD-PH and PD-nPH (permutation p-value=1). The average duration of the poke duration for PD-PH was 0.75±5.24 seconds (mean ± SD) and 0.73±2.82 seconds (mean ± SD) for PD-nPH (Fig.S2A-B). Spatial analysis of the movement revealed no difference in the distance between the pokes between PD-PH and PD-nPH (permutation p-value = 0.3). Average surface explication for PD-PH was 17.83±18.4 mm (mean ± SD) and 23.89±21.05 mm for PD-nPH (mean ± SD)

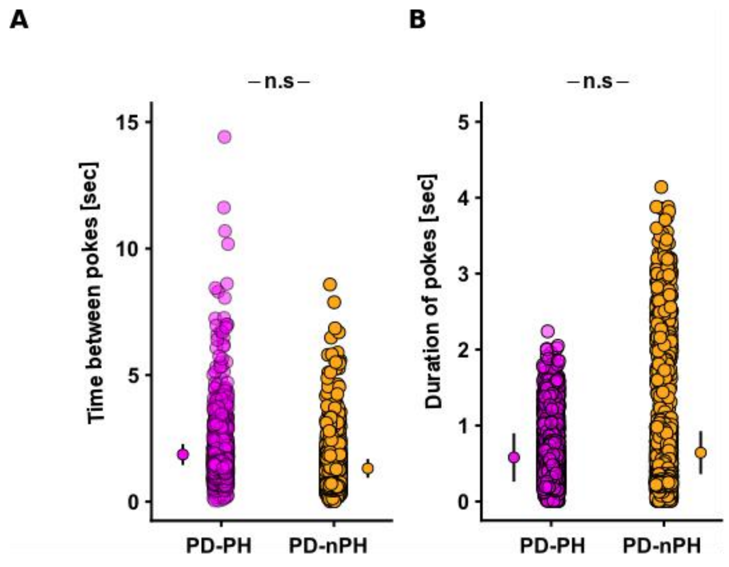


**Figure S2. Analysis of the movement patterns during the sensorimotor stimulation.** A. Mixed effects linear regression between the time between pokes for PD-PH (purple) and PD-nPH (yellow). The duration did not differ significantly (permutation p-value=0.29) in the time between two pokes (inter-pokes-interval). B. Mixed effects linear regression between duration of the pokes. The duration of the pokes did not differ significantly (p-value=1). Error bar represent 95% confidence interval.

PD-PH were not significantly slower in performing poking movement than HC (1.57±2.08; mean ± SD) (permutation p-value=0.097). The duration of each poke did not differ between PD-PH and HC (permutation p-value=0.076), the average duration of the poke duration for HC was 0.49±0.39 seconds (mean ± SD). No differences were observed in the spatial aspects of the movement between PD-PH and HC (permutation p-value = 0.067). The average surface explored for HC was 29.45±25.78 mm (mean ± SD).

These movement data show that both PD groups and the elderly HC were well able to carry out sensorimotor stimulation during the robotic procedure and, importantly, that movement patterns did not differ between both patient groups.

### *Supplementary S14*

#### *riPH are not due to clinical differences between PD-PH and PD-nPH*

All patients were treated with anti-parkinsonian medications, but there was no significant difference in medication between both patient groups (Table S1). Although clinical experience and research has associated hallucinations with dopaminergic treatment[11,14], the exact role between the dopaminergic system and hallucinations is currently debated[15,16]. Thus, PD patients can experience PH before starting any dopaminergic medication [17] and the use of levodopa and dopamine agonists was not found to modulate occurrence of PH[15]. Motor fluctuations have also been linked to PH [18] (for a review [15]) and it is known that PH and other hallucinations occur more frequently in advanced stages of PD [16]. Thus, the difference in riPH between the two PD groups was not related to differences in disease duration, motor impairment, motor complications, or to dopaminergic treatment or hyperdopaminergic behavior (no significant differences between PD-PH and PD-nPH: all p-values>0.05; two-tailed permutation test; Table S1-2). Analyzing several other clinical and demographic variables (including motor impairment, dopaminergic treatment, and) that have been associated with symptomatic hallucinations in PD (e.g. [18]). We found no evidence for a difference between the two sub-groups of patients (all p-values>0.05; Table S1 for demographic of sub-groups).

Between the two sub-groups of patients with PD, there were no statistically significant differences (p-values>0.05) in the performance on the Montreal Cognitive Assessment (MoCA), disease duration, dopaminergic treatments (levodopa daily equivalent dosage), apathy, hyperdopaminergic behavior (QUIP-RS), motor impairment (MDS-UPDRS-III) and motor complications (MDS-UPDRS-IV). Patients that felt the presence as a symptom of the disease, PD-PH (vs. PD-nPH), had a higher score on the score for risk of psychosis (PQ-16, part 1- assessing hallucinations, but not for part 2 - assessing the level of distress associated with the occurrence of hallucinations).

Collectively, these results suggest that the differences in robot induced-PH between PD-PH and PD-nPH cannot be explained by differences in the degree of neurodegeneration, dopaminergic treatment, or motor fluctuations (or any of the other clinical variables we measured).

### Study 2

#### Study 2.1: riPH are associated with activation of a subcortical-cortical sensorimotor network in healthy subjects

***Supplementary S15***

***Robotic system***

The MR-compatible robotic system used to generate the sensorimotor conflicts was composed of a front and a back robot (Fig.2A in the main text;[19]). The front-robot of the MR-compatible robot contained a carbon-fiber rod attached to a slider allowing the participant to move along two directions (Fig.2A in the main text) and measured the movements. Movements of the carbon-fiber rod were electronically translated into movements of the back-robot. The back-robot was composed of a roller that touched the participant's back with stroking and tapping movements (for general performance of the robotic system see [19]). The back-robot's shape was adapted to the spatial dimensions of the scanner bore and a wooden mattress structure with a central slit was designed to allow the contact part of the back-robot to touch the back of the participants. The performance of the robotic system was previously validated inside a 3T and 7T MR scanner with a phantom[19]. Visual Studio 2013 interface (Microsoft) was used to control the robotic system.

The robotic system used in this study differed from the one used in the study 1 and of Blanke and colleagues[9] in multiple aspects. First, the participants were in the supine position compared to the standing position. Secondly, due to the spatial constrains of the MR-environment, the movement of the participants were limited to the middle back and not the whole back and participants had less degree of freedom: they could only move in X (along the body) and Z (towards the body) directions. All these different aspects might have led to the decrease of intensity of the PH induction compared to the standing robot used in the previous study[9].

### *Supplementary S16*

#### *Mock scanner: pilot study*

Here, we tested whether we could induce PH in supine position in a mock scanner using the MRI robot (i.e. riPH). All participants (n=24; 16 women, mean age±SD: 24.6±2.8 years old) had no history of neurological or psychiatric disorders. All participants were right handed as assessed by the Edinburg Handedness Inventory [20] (mean index: 81.0± 16.3 (SD) and range: 40-100). All participants provided written informed consent prior to the experiment. The study was approved by the Cantonal Ethics Committee of Geneva (Commission Cantonale d'Ethique de la Recherche-CCER). The Mock scanner (MRI Simulator, Psychology Software Tools, Inc.) mimicked the scanner environment as well as the noise of the echo-planar imaging sequence. Participants were asked to perform repetitive movements with their right hand and this operated the front-robot, the movements of which were translated to the back-robot that provided tactile feedback to our participants' backs. In two conditions, tactile feedback was delivered either synchronously with the participants' movements (synchronous control condition, sync) or with a delay (asynchronous condition, async) that was previously shown to induce the PH in healthy participants. In a third condition (desynchronous condition, desync), movements of the back- robot consisted of a pre-recorded sequence. Each condition lasted for 3 minutes, was repeated once, and given in random order. After each condition, a questionnaire adapted from [9] was filled where participants were asked to rate their degree of agreement or disagreement on a Likert scale from 0 to 6.

### *Supplementary S17*

#### *Mock Scanner (pilot study): Questionnaire results*

All the ratings are summarized in Table S5.

*Passivity experience ("I felt as if someone else was touching my body")*

In line with prior work[21], we found that asynchronous robotic stimulations were associated with higher passivity experience than synchronous stimulation (main effect of Synchrony: permutation p-value < 0.001) with higher ratings in the async and desync conditions compared to the sync condition (post-hoc test: t(46) = 2.16, p-value = 0.035 and t(46) = 4.75, p-value < 0.001, respectively) and higher ratings in the desync compared to the async condition (post-hoc test: t(46) = 2.59, p-value = 0.012). These results further indicate a difference in the passivity experience between the desync and the async condition. This can be explained by the fact that the desync condition is a pre-recorded movement sequence played on the back of the participants which is being completely decoupled with the participant's movement.

*PH ("I felt as if a presence or someone was behind me")*

A main effect of Synchrony was also found for PH (permutation p-value < 0.001) with higher ratings in the desync and async condition compared to the sync condition post-hoc test: (t(46) = 4.14, p-value < 0.001 and t(46) = 2.92, p-value = 0.00053, respectively). No significant difference between the async and desync ratings was found (post-hoc test: t(46) = 1.13, p-value = 0.26). Contrary to passivity experience, the desync condition did not elicit significantly higher ratings than the async condition suggesting that both condition generating sensorimotor conflicts can equally induce PH. Taken together, these results confirm previous findings in showing that passivity experience and PH are induced in the presence of strong sensorimotor conflicts, in line with the results found by Blanke and colleagues [9].

*Self-touch ("I felt as if I was touching my body")*

Regarding self-touch, only a trend for a main effect of Synchrony was found (permutation p-value=0.062), with a tendency for higher ratings in the sync condition compared to async and desync (Table S5 for ratings).

| Question | Synchrony | Mean | Standard deviation |
|---|---|---|---|
| Self-touch | Desync | 1.58 | 1.84 |
| Self-touch | Async | 2.08 | 2.10 |
| Self-touch | Sync | 2.54 | 2.48 |
| I felt as if I was touching someone else's body | Desync | 0.71 | 1.55 |
| I felt as if I was touching someone else's body | Async | 0.75 | 1.62 |
| I felt as if I was touching someone else's body | Sync | 0.50 | 1.29 |
| Passivity experience | Desync | 3.96 | 1.97 |
| Passivity experience | Async | 2.71 | 2.14 |
| Passivity experience | Sync | 1.67 | 1.97 |
| PH | Desync | 2.08 | 1.82 |
| PH | Async | 1.67 | 1.81 |
| PH | Sync | 0.67 | 1.55 |
| Control (I felt as if I had no body) | Desync | 0.67 | 1.09 |
| Control (I felt as if I had no body) | Async | 0.42 | 0.88 |
| Control (I felt as if I had no body) | Sync | 0.33 | 0.64 |
| Control (I felt as if I had two right hands) | Desync | 0.83 | 1.40 |
| Control (I felt as if I had two right hands) | Async | 0.75 | 1.51 |
| Control (I felt as if I had two right hands) | Sync | 0.63 | 1.06 |

**Table S5. Mean ratings for all questions used in the mock scanner study**

*Movement analysis*

To ensure that riPH were not due to any movement differences across experimental conditions, we calculated the total distance that each participant moved the front-robot. Analysis revealed no significant difference between the total distance covered in the synchronous versus asynchronous condition (permutation p-value = 0.96).

### *Supplementary S18*

#### *fMRI behavioral study 2.1: Questionnaire results*

The questionnaire included only the first six questions of the mock scanner pilot study: "I felt as if I was touching my body", "I felt as if I was touching someone else's body", " I felt as if I had no body", "I felt as if I had two right hands", "I felt as if someone else was touching my body" and "I felt as if a presence or someone was behind me".

*Passivity experience*

Participants reported stronger passivity experiences in the asynchronous condition compared to the synchronous condition (main effect of Synchrony, permutation p-value < 0.001; Table S6).

| Question | Synchrony | Mean | Standard deviation |
|---|---|---|---|
| Self-touch | Async | 3.28 | 2.25 |
| Self-touch | Sync | 3.72 | 2.23 |
| I felt as if I was touching someone else's body | Async | 1.12 | 1.81 |
| I felt as if I was touching someone else's body | Sync | 0.88 | 1.59 |
| Passivity experience | Async | 3.40 | 2.25 |
| Passivity experience | Sync | 2.08 | 2.14 |
| PH | Async | 1.68 | 1.86 |
| PH | Sync | 1.04 | 1.65 |
| Control (I felt as if I had no body) | Async | 1.04 | 1.62 |
| Control (I felt as if I had no body) | Sync | 0.80 | 1.76 |
| Control (I felt as if I had two right hands) | Async | 0.56 | 1.33 |
| Control (I felt as if I had two right hands) | Sync | 0.36 | 0.76 |

**Table S6. Mean ratings for all questions of the fMRI questionnaire**

### *Supplementary S19*

#### *riPH-network in healthy subjects*

We also analyzed fMRI data recorded in two control conditions that allowed us to control for two aspects of sensorimotor stimulation that are not related to PH and determined the brain regions that were commonly activated by either of the sensorimotor conditions (synchronous, asynchronous; Fig.S3A-B) vs. the control conditions (motor, touch; Fig.S3C-D). In the motor control condition, participants were asked to repeatedly move the front-robot with their right hand but did not receive any tactile feedback on their back (Fig.S3C). In the touch control condition, participants received touch feedback on their backs, but were not performing any movement with their right hand (the back-robot was actuated by a previously recorded movement sequence) (Fig.S3D).

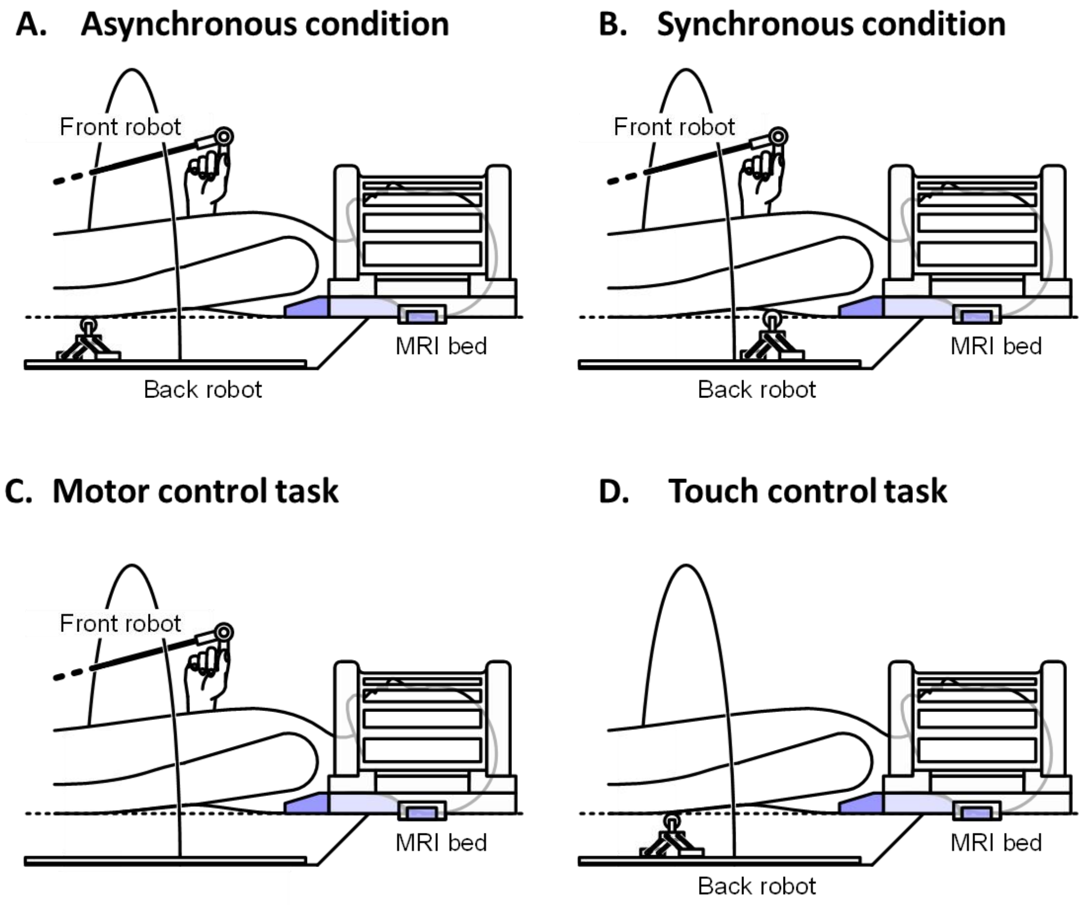


**Figure S3. The different conditions assessed with MR-compatible robotic system**

The MR robotic system consisted of two parts: a front robot composed by a carbon fibre rod with which the participants performed the movement in 2 directions (X and Z) and a back robot that reproduced the movement of the front robot in the back of the participants. Different conditions were tested: (A) an asynchronous condition where the back robot was delayed of 500 ms compared to the front robot, (B) a synchronous condition in which no delay was introduced between the front robot and the back robot. In addition, in the fMRI study, two conditions were added: a motor control task, in which the participant was just performing the movements without any tactile feedback on the back (C) and a touch control task in which the participant only received the tactile feedback on the back without any movement (D). The contact part is composed of a roller effector that enables to touch the back of the participant. Two ultrasonic motors (USR60-E3NT, Shinsei) enable the effector to move. A home-made mattress was designed with an aperture to allow robotic stroking on the participant's back, while lying down.

### *Supplementary S20*

#### *riPH are associated with activation of two sensorimotor networks in healthy subjects*

Fig.S4A shows the activations when comparing the asynchronous condition with the motor plus touch control condition, revealing a large cortical-subcortical network including the left sensorimotor cortex (including adjacent parts of premotor cortex and superior parietal cortex), bilateral SMA and adjacent parts of cingulate cortex, bilateral putamen, the right ventral premotor cortex, the right inferior parietal cortex (IPL) and the right cerebellum (Table S8). Similar regions were found for the contrast between the synchronous and the motor plus touch control condition (Fig.S4B; Table S8). The synchronous versus asynchronous contrast did not show any significant brain activations. We also correlated riPH ratings or passivity experiences with brain regions activated more during the asynchronous condition compared to the synchronous condition and found no significant correlation (all p-values>0.05 after correcting for multiple comparisons). Activations from the conjunction analysis (Fig.2E main text) also did not correlate with PH ratings or passivity experiences (all p-values>0.05 after correcting for multiple comparisons).

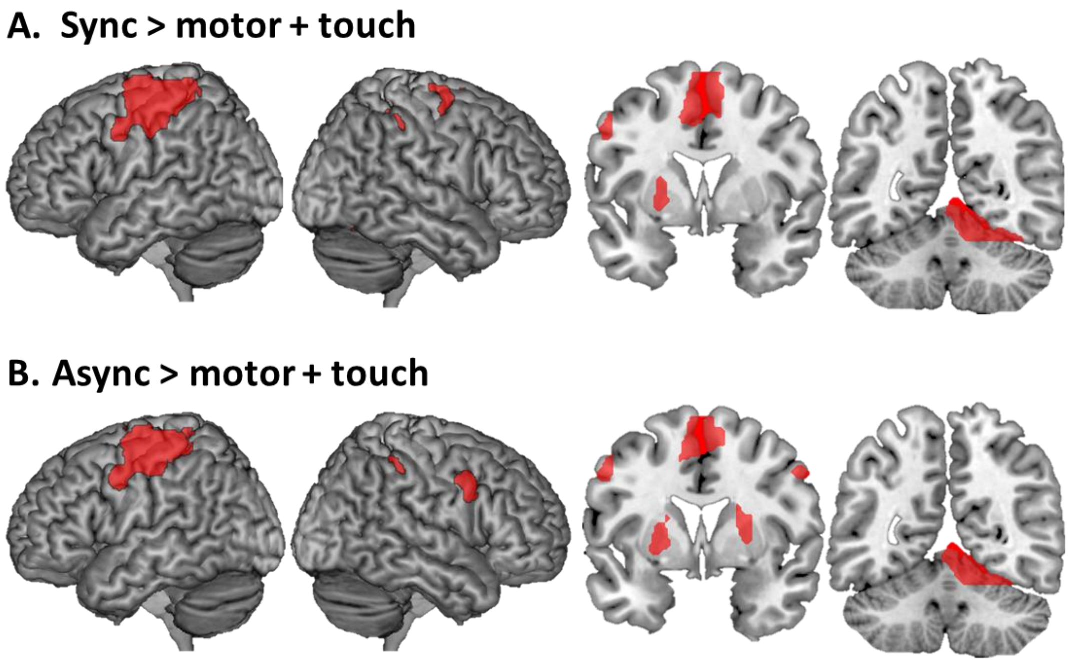


**Figure S4. Robot-induced PH network. A.** Brain regions responding to spatial sensorimotor conflict between the right-hand movement and the feedback on the back of the participants. **B.** Brain regions reflecting the spatio-temporal sensorimotor conflict.

| Regions | Voxels | BA | MNI x | MNI y | MNI z | Peak level t value | Cluster-level *p* value FWE |
|---|---|---|---|---|---|---|---|
| R. medial prefrontal cortex (mPFC) | 770 | 8/9/32 | 5 | 41 | 47 | 6.2 | 0.001 |
| R. ventral premotor cortex (vPMC)/ Inferior Frontal Gyrus (Opercularis and triangularis) (IFG) | 708 | 45/48 | 51 | 18 | 29 | 5.53 | 0.001 |
| R. Anterior Insula (Ins) | 566 | 47 | 36 | 24 | -2 | 4.78 | 0.004 |
| R. posterior Middle Temporal Gyrus (pMTG) | 479 | 37 | 54 | -54 | 0 | 4.99 | 0.01 |

**Table S7. Spatio-temporal sensorimotor conflict PH regions.** Regions activated during the contrast asynchronous > synchronous.

| Regions | Voxels | BA | MNI coordinates x | y | z | Peak level t value | Cluster-level p value FWE |
|---|---|---|---|---|---|---|---|
| ***Asynchronous > motor + touch*** | | | | | | | |
| L. Sensorimotor cortex (primary motor cortex (M1), primary somatosensory cortex (SI), supplemental motor area (SMA), middle cingulate cortex (MCC), Superior parietal lobe (SPL)) | 9894 | 2/3/4/6/40 | -26 | -16 | 58 | 7.93 | p<0.001 |
| R. Cerebellum | 2840 | | 11 | -58 | -14 | 7.99 | p<0.001 |
| R. Putamen / Globus pallidum | 599 | | 22 | 3 | 8 | 6.33 | p<0.01 |
| L. Putamen / Globus pallidum | 560 | | -22 | 1 | 5 | 6.07 | p<0.01 |
| R. Inferior parietal lobe/supramarginal gyrus (SMG) | 503 | 2/40 | 41 | -37 | 47 | 4.1 | p<0.01 |
| R. ventral premotor cortex | 357 | 6 | 55 | 8 | 38 | 5.58 | p<0.05 |
| ***Synchronous > motor + touch*** | | | | | | | |
| L. Sensorimotor cortex (primary motor cortex (M1), primary somatosensory cortex (SI), supplemental motor area (SMA), middle cingulate cortex (MCC), Superior parietal lobe (SPL)) | 12843 | 2/3/4/6/40 | -25 | -18 | 57 | 8.85 | p<0.001 |
| R. Cerebellum | 3057 | | 12 | -57 | -14 | 8.17 | p<0.001 |
| R. Inferior parietal lobe/supramarginal gyrus (SMG) | 600 | 2/40 | 40 | -34 | 45 | 4.8 | p<0.01 |
| L. Putamen / Globus pallidum | 449 | | -22 | 0 | 5 | 7.34 | p<0.05 |
| R. Superior frontal gyrus / dorsal premotor cortex | 385 | 6 | 28 | -8 | 65 | 4.62 | p<0.05 |
| ***Conjunction between the asynchronous > motor + touch and synchronous > motor + touch*** | | | | | | | |
| L. Sensorimotor cortex (primary motor cortex (M1), primary somatosensory cortex (SI), middle cingulate cortex (MCC), Superior parietal lobe (SPL)) | 12026 | 2/3/4/6/40 | -26 | -18 | 57 | 9.99 | p<0.001 |
| Supplemental motor area (SMA) | | 6 | -5 | -6 | 56 | 8.32 | p<0.001 |
| R. Cerebellum | 2687 | | 11 | -57 | -14 | 9 | p<0.001 |
| R. Inferior parietal lobe/supramarginal gyrus (SMG) | 593 | 2/40 | 40 | -34 | 45 | 4.59 | p<0.01 |
| L. Putamen / Globus pallidum | 517 | | -23 | 0 | 4 | 6.44 | p<0.01 |

**Table S8. Robotically induced brain activations.**

### Study 2.2: Common PH-network for sPH and riPH

### *Supplementary S21*

#### *Lesion network mapping analysis*

In order to assess the functional network derived from PH, we applied lesion network mapping [22]. This method has the advantage of not requiring functional neuroimaging data from patients and of accounting for the possibility that symptoms may arise from remote brain regions connected to the lesioned brain region rather than the damaged area itself [23,24]. The PH-lesions reported by Blanke and colleagues [21] were used as seed ROIs except one lesion which was covering the whole brain, resulting in eleven lesions for the analysis.

*fMRI acquisition*

Resting state and T1-weighted structural data from 151 healthy participants obtained from the publicly available Enhanced Nathan Kline Institute Rockland Sample[25] was used. All participants were right-handed and aged between 19 to 40 years (25.8 ± 5.5 years, 83 females). Scans were acquired with a 3T Siemens Magneton TrioTim syngo. For the resting state data, a multiband EPI sequence was used (multiband factor = 4, 64 continuous slices, TR = 1.4 s, TE = 30 ms, filp angle = 65°, slice thickness = 2 mm) and 404 scans were collected. For each participant, an anatomical image was recorded using a T1-weighted MPRAGE sequence (TR = 1.9 s, TE = 2.52 ms, Inversion time = 900 ms, flip angle = 9°, 1 mm isotropic voxels, 176 slices per slab and FOV = 250 mm).

*Data analysis*

The pre-processing steps were performed using SPM12 toolbox (Wellcome Departement of Cognitive Neurology, Institute of Neurology, UCL, London, UK) in Matlab (R2016b, Mathworks). The first four functional scans were discarded from the analysis to allow for magnetic saturation effects: the analysis was performed on the 400 remaining scans. Slice timing correction and spatial realignment was applied to individual functional images. The anatomical image was then co-registered with the mean functional image and segmented into grey matter, white matter and cerebro-spinal fluid tissue. The functional and anatomical scans were then normalized to the Montreal Neurological Institute space (MNI space). Finally, the functional scans were spatially smoothed with a 5 mm full-width at half-maximum isotropic Gaussian kernel.

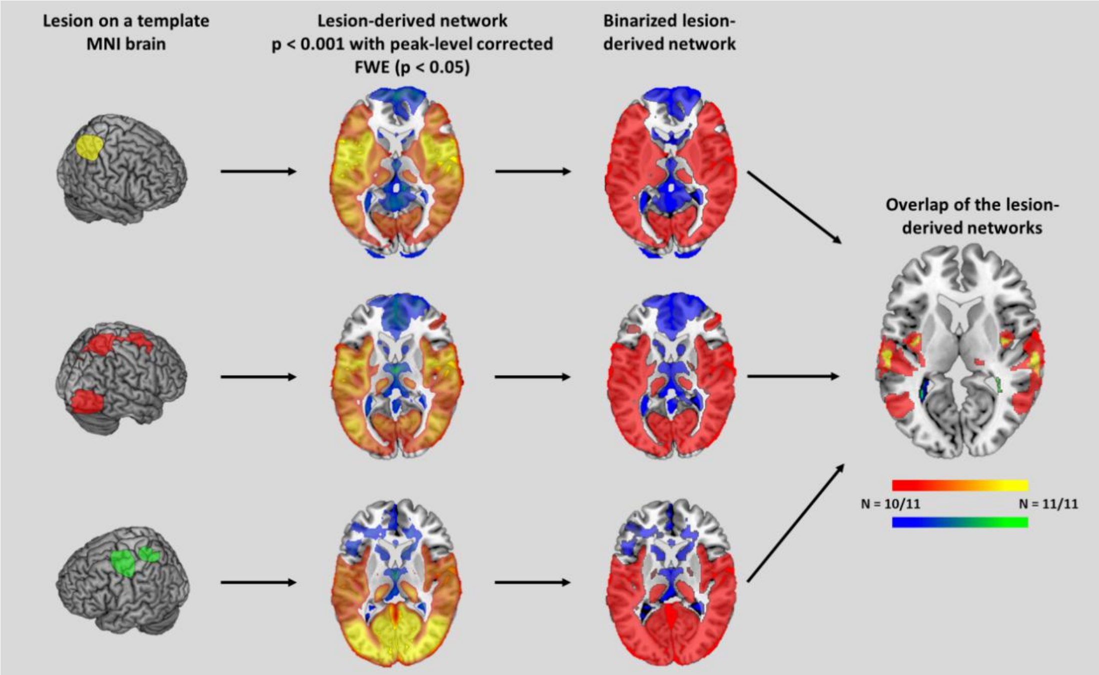


**Figure S5. Lesion network mapping analysis.**

The steps of the lesion network mapping analysis are shown: first the lesion is mapped to a template brain, then this lesion is used as a seed ROI in a resting state functional analysis performed on a normative database. The network obtained for each lesion is thresholded at $p < 0.001$ with peak level corrected FWE ($p < 0.05$). All the lesions-derived networks are binarized and overlap to identify the regions functionally connected to most of the lesions.

### *Supplementary S22*

#### *Lesion network mapping: control analysis*

To exclude that these regions are involved in hallucinations more generally, the same method was applied to a control group of eleven patients suffering from structured visual hallucination (VH)[26]. The sPH-network was defined as those PH regions that were not overlapping with the visual hallucination derived network.

In addition, we determine whether the riPH-network was specifically connected to the lesions causing PH as opposed to the lesions causing VH. Therefore, for each of the 126 subjects in the database, the regionally-averaged resting-state BOLD signal time courses were extracted from each PH and VH lesion and riPH-network (Fig.2D-E in the main text) and were pairwise correlated (Fisher Z-transformed Pearson correlation) to establish the functional connectivity matrix. For each lesion location, we averaged the connectivity measures for the riPH-networks. Then, we compared statistically the connectivity between the two groups (PH vs. VH) using two sample t-test.

### *Supplementary S23*

#### *Results*

#### *Lesion network mapping control analysis*

To exclude that these regions are involved in hallucinations more generally, the same method was applied to a control group of eleven patients suffering from structured visual hallucination (VH)[26], revealing a VH derived network consisting of mostly distinct regions (in bilateral TPJ, dorsal premotor cortex (dPMC), the left middle and superior occipital cortex, left thalamus and hippocampus (Table S11), as well as three common regions (bilateral posterior to middle STG and adjacent parts of parietal cortex; left PMC). Further analysis showed that the brain lesions causing sPH were more strongly connected with the riPH-network (as defined in healthy participants; study2.1) than the lesions causing visual hallucinations (difference between the two groups of lesions: $t(18)=2.74$, p-value=0.013, Fig.S6). A sPH-network was defined as those PH regions that were not overlapping with the visual hallucination derived network.

| Regions | Overlap | Hemisphere | Voxels | BA | MNI coordinates | | |
|---|---|---|---|---|---|---|---|
| | | | | | x | y | z |
| ***Positive correlation*** | | | | | | | |
| Superior temporal gyrus (STG) | 11 | Right | 770 | 22 | 62 | 25 | 13 |
| | 11 | Left | 582 | 22 | -58 | -29 | 13 |
| Insula | 11 | Right | 124 | 48 | 37 | -6 | 12 |
| | 11 | Left | 135 | 48 | -37 | -7 | 9 |
| | 11 | Left | 81 | 48 | -35 | -9 | -8 |
| Postcentral sulcus | 11 | Left | 111 | 48 | -58 | -16 | 19 |
| Middle cingulate cortex (MCC) | 11 | Right | 53 | | 9 | -11 | 38 |
| | 11 | Left | 100 | | -9 | -11 | 37 |
| Inferior frontal operculum/ ventral premotor cortex (vPMC) | 11 | Right | 86 | 45/48 | 42 | 17 | 23 |
| Temporo-parietal junction (TPJ): STG, MTG (only right), supramarginal gyrus (SMG), rolandic operculum, vPMC | 10 | Right | 7153 | 21/22/48 | 56 | -18 | 18 |
| | 10 | Left | 5318 | 21/22/48 | -52 | -16 | 16 |
| Fusiform area | 10 | Right | 2842 | 19/37 | 37 | -52 | -16 |
| | 10 | Left | 2916 | 19/37 | -36 | -53 | -15 |
| Middle temporal gyrus (MTG) | 10 | Left | 1292 | 37 | -48 | -62 | 11 |
| Dorsal premotor cortex (dPMC) | 10 | Right | 370 | 6 | 44 | -5 | 53 |
| | 10 | Left | 308 | 6 | -40 | -8 | 51 |
| Amygdala | 10 | Right | 295 | 36 | 29 | 3 | -24 |
| | 10 | Left | 112 | 36 | -26 | 2 | -26 |
| Thalamus | 10 | Right | 126 | | 15 | -25 | 2 |
| | 10 | Left | 120 | | -12 | -27 | -2 |
| Cerebellum | 10 | Left | 107 | | -10 | -65 | -46 |
| Hippocampus | 10 | Rigth | 70 | | 23 | -36 | -2 |
| | 10 | Left | 90 | | -20 | -37 | -1 |
| Putamen | 10 | Right | 69 | | 36 | -10 | -8 |
| Cuneus | 10 | Right | 68 | 18 | 17 | -70 | 26 |
| | 10 | Left | 68 | 18 | -14 | -72 | 22 |
| Suppplemental motor area (SMA)/Superior frontal gyrus | 10 | Left | 58 | 6 | -18 | -8 | 68 |
| ***Negative correlation*** | | | | | | | |
| Caudate | 10 | Right | 70 | | 17 | -13 | 27 |

**Table S9. Symptomatic PH-derived network**. Brain areas that showed positive and negative correlation with most of the lesions (100% or 90% of overlap). Regions in the white matter were not reported.

| Regions | Overlap | Hemisphere | Voxels | BA | MNI coordinates | | |
|---|---|---|---|---|---|---|---|
| | | | | | x | y | z |
| ***Positive correlation*** | | | | | | | |
| Superior temporal cortex (TPJ) | 10 | Right | 734 | 22/48 | 60 | -14 | 9 |
| | 10 | Left | 148 | 42/22 | -61 | -32 | 14 |
| | 10 | Left | 147 | 22 | -59 | -9 | -8 |
| | 10 | Left | 92 | 48 | -51 | -21 | 5 |
| Middle and superior occipital cortex/ Inferior parietal lobule | 10 | Left | 326 | 39/19 | -38 | -70 | 28 |
| Hippocampus/parahippocampus | 10 | Left | 118 | 20 | -27 | -31 | -14 |
| Thalamus/lingual area | 10 | Left | 108 | 27 | -14 | -30 | -2 |
| Precentral cortex (dPMC) | 10 | Right | 74 | 6 | 53 | -3 | 44 |
| | 10 | Left | 51 | 6 | -45 | -7 | 51 |

**Table S10. VH-derived network.** Brain areas that showed positive correlation with 90 % of the VH lesion locations (only the regions in the grey matter are reported). There was no overlap for all the lesions.

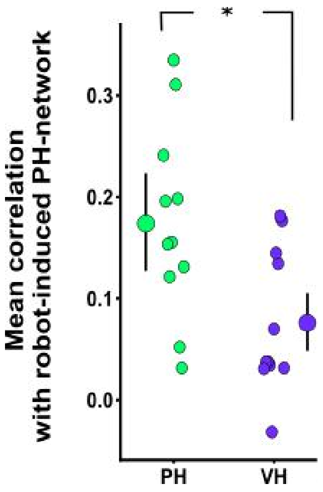


**Figure S6. Lesion connectivity with the robot-induced PH-network.** Lesions causing PH had greater functional connectivity with the riPH-network compared to VH lesions. * p-value<0.05.

### *Supplementary S24*

#### *Common PH-network*

The common PH-network (cPH-network) was composed of regions overlapping between the sPH-network and the riPH-network and consisted in three anatomical regions in right IFG, right pMTG and two almost continuous PMC clusters (considered as one ROI for the following analysis).

### Study 3

#### Study 3.1: Disrupted functional connectivity in cPH-network accounts for sPH in patients with Parkinson's disease

### *Supplementary S25*

#### *Participants*

Data from thirty participants were analyzed in this study. All patients were prospectively recruited from a sample of outpatients regularly attending to the Movement Disorders Clinic at Hospital de la Santa Creu i Sant Pau (Barcelona) based on the fulfilling of MDS new criteria for PD with minor hallucinations (PD-PH) — sense of presence and/or passage hallucinations (n=15) — and without hallucinations (PD-nPH; n=15). Informed consent to participate in the study was obtained from all participants. The study was approved by the local ethics committee. The same dataset has been previously used in [27]. Patients were diagnosed by a neurologist with expertise in movement disorders. Each patient was interviewed regarding years of formal education, disease onset, medication history, current medications, and dosage (levodopa daily dose). Motor status and stage of illness were assessed by the MDS-UPDRS-III[28]. The PD-PH and PD-nPH groups did not differ for age, disease duration, dopaminergic doses, motor severity, cognition, depression, anxiety, and apathy (Table S11). All participants were on stable doses of dopaminergic drugs during the 4 weeks before inclusion. Patients were included if the hallucinations remained stable during the 3 months before inclusion in the study. No participant had used or was using antipsychotic medication.

| | **PD-PH (N = 15)** | **PD-nPH (N = 15)** | ***p-values*** |
|---|---|---|---|
| Age | 70.9 ± 1.5 | 65.9 ± 1.94 | 0.06 |
| Gender | 9 (M) | 10 (M) | 0.7 ($\chi$2) |
| MoCA | 25.3 ± 0.8 | 24 ± 1 | 0.3 |
| PD-CRS | 91.5 ± 4 | 94.2 ± 4.1 | 0.67 |
| PD-CRS (frontal) | 62.9 ± 3.8 | 65.7 ± 3.9 | 0.62 |
| PD-CRS (posterior) | 28.7 ± 0.4 | 28.5 ± 0.4 | 0.83 |
| UPDRS III | 21.7 ± 2.4 | 25.3 ± 2.03 | 0.2 |
| LEDD (mg/day) | 722.1 ± 73.8 | 581 ± 80.2 | 0.2 |
| Dopamine agonists (mg/day) | 151.3 ± 31.7 | 151.3 ± 31.7 | 0.9 |
| Disease Duration (years) | 5.3 ± 0.9 | 3.7 ± 0.6 | 0.2 |

**Table S11. Clinical variables.**

Exclusion criteria were history of major psychiatric disorders, cerebrovascular disease, conditions known to impair mental status other than PD, and the presence of factors that prevented MRI scanning (e.g. claustrophobia, MRI non-compatible prosthesis). Patients with focal abnormalities in MRI or non-compensated systemic diseases (e.g. diabetes, hypertension) were also excluded. In patients with motor fluctuations, cognition was examined during the "on" state. All participants were on stable doses of dopaminergic drugs during the 4 weeks before inclusion. Patients were included if the hallucinations remained stable during the 3 months before inclusion in the study. No participant had used or was using antipsychotic medication. All subjects had normal or corrected-to-normal vision. Informed consent to participate in the study was obtained from all participants. The study was approved by the local ethics committee.

Presence and type of minor hallucinations was assessed using the Hallucinations and Psychosis item of the MDS-UPDRS Part I. Participants with a sense of presence and/or passage hallucinations at least weekly during the last month were categorized as minor hallucinations. Cognitive functions were assessed by the Parkinson's Disease-Cognitive Rating Scale (PD-CRS)[29]. Apathy was assessed with the Starkstein Apathy Scale[5].

We favored the analysis of resting state fMRI over performing the robotic stimulation within the MRI, because for PD patient performing long motor task (as required by the MRI to have a good signal to noise ratio) can be particularly tiring, and therefore exacerbating the tremor. Thus, the probability to have poor data quality and a high rate of patient willing to interrupt the experiment prematurely was too high.

### *Supplementary S26*

#### *Image acquisition & Image processing*

MRI scans were acquired with a 3T Philips Achieva. T1 weighted scans were obtained using a Magnetization Prepared Rapid Acquisition Gradient Echo (MPRAGE) sequence (TR = 500 ms, TE = 50 ms, flip angle = 8, field of view [FOV] = 23 cm with in-plane resolution of 256 × 256 and 1mm slice thickness). Resting-state functional MRI images were collected using an 8-minute sequence (TR = 2000 ms, TE = 30 ms, flip angle = 78, FOV = 240 mm, slice thickness = 3 mm).

Data analysis and standard pre-processing was performed using the functional connectivity toolbox CONN (https://www.nitrc.org/projects/conn/) and Statistical Parametrical Mapping (SPM 12) (http://www.fil. ion.ucl.ac.uk/spm/) for Matlab. Functional images were corrected for slice time and motion, co-registered with a high-resolution anatomical scan, normalised into Montreal Neurological Institute

(MNI) space, resampled to 2 x 2 x 2 mm3, and smoothed with an 8 mm3 full width at half maximum (FWHM) Gaussian kernel for each subject. To estimate the excessive movement, the mean frame-wise displacement (FD) during the scanning was estimated with the exclusion threshold of 0.5 mm. The groups did not differ by the movement over the scanning period ($t = 1.18$, $p = 0.12$ with the mean FD of 0.29 ± 0.15 mm and 0.23 ± 0.16 mm for PD-PH and PD-nPH groups respectively) and did not reach the excessive movement threshold. Following the standard pipeline for confound removal of the CONN toolbox, the individual time courses of the segmented white matter and cerebrospinal fluid, the 6 motion parameters with rigid body transformations and their first-order derivatives, and global signal time courses were extracted and regressed out of the data. Regressions were performed for the entire time-series. The blood oxygenation level dependent (BOLD) signal data were passed through a band filter of 0.01-0.1 Hz. A whole-brain grey matter mask in MNI space restricted data analysis.

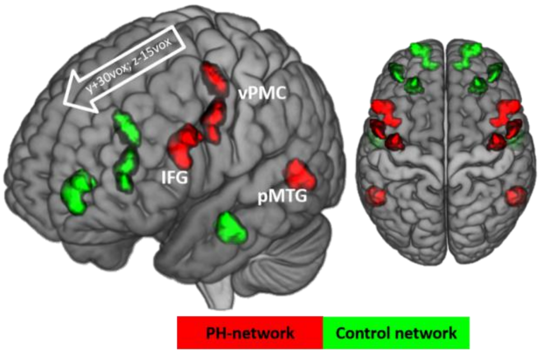


**Figure S7. Control regions for the resting state fMRI analysis of PD patients.** Bilateral PH-network areas (red) shifted forward (green): Inferior frontal gyrus (IFG) x±20 y+30 z-15; ventral premotor cortex (vPMC) x±10 y+30 z-15; posterior middle temporal gyrus (pMTG) x y+30 z-15.

| Connections | Variable Importance |
|---|---|
| Left-IFG - Left pMTG | 1 |
| Left pMTG - Right vPMC | 0.913 |
| Left IFGlh - Right vPMC | 0.797 |
| Left pMTG - Right pMTG | 0.710 |
| Left IFG - Right pMTG | 0.652 |
| Left IFGlh - Left vPMC | 0.348 |
| Right IFG - Right vPMC | 0.333 |
| Right pMTG - Right vPMC | 0.333 |
| Left pMTG - Left vPMC | 0.319 |
| Right IFG - Right pMTG | 0.225 |
| Right pMTG - Left vPMClh | 0.203 |
| Left vPMC - Right vPMC | 0.188 |
| Left IFG - Right IFG | 0.174 |
| Right IFG - Left vPMC | 0.123 |
| Right IFG - Left pMTG | 0 |

**Table S12. Contribution of each connectivity to the classification of PD-PH, values scaled between zero and one.**

*Supplementary References*


1. Nasreddine, Z. S. *et al.* The Montreal Cognitive Assessment, MoCA: a brief screening tool for mild cognitive impairment. *J Am Geriatr Soc* **53**, 695–699 (2005).

2. Carson, N., Leach, L. & Murphy, K. J. A re-examination of Montreal Cognitive Assessment (MoCA) cutoff scores. *International Journal of Geriatric Psychiatry* **33**, 379–388 (2018).

3. Tomlinson, C. L. *et al.* Systematic review of levodopa dose equivalency reporting in Parkinson's disease. *Movement Disorders* **25**, 2649–2653 (2010).

4. Weintraub, D. *et al.* Questionnaire for Impulsive-Compulsive Disorders in Parkinson's Disease-Rating Scale. *Mov. Disord.* **27**, 242–247 (2012).

5. Starkstein, S. E. *et al.* Reliability, validity, and clinical correlates of apathy in Parkinson's disease. *J Neuropsychiatry Clin Neurosci* **4**, 134–139 (1992).

6. Loewy, R. L., Bearden, C. E., Johnson, J. K., Raine, A. & Cannon, T. D. The prodromal questionnaire (PQ): preliminary validation of a self-report screening measure for prodromal and psychotic syndromes. *Schizophr. Res.* **77**, 141–149 (2005).

7. de Chazeron, I. *et al.* Validation of a Psycho-Sensory hAllucinations Scale (PSAS) in schizophrenia and Parkinson's disease. *Schizophr. Res.* **161**, 269–276 (2015).

8. Cilia, R. *et al.* Dopamine dysregulation syndrome in Parkinson's disease: from clinical and neuropsychological characterisation to management and long-term outcome. *J. Neurol. Neurosurg. Psychiatry* **85**, 311–318 (2014).

9. Blanke, O. *et al.* Report Neurological and Robot-Controlled Induction of an Apparition. *Current Biology* **24**, 2681–2686 (2014).

10. Hara, M. *et al.* A novel approach to the manipulation of body-parts ownership using a bilateral master-slave system. in *2011 IEEE/RSJ International Conference on Intelligent Robots and Systems* 4664–4669 (2011). doi:10.1109/IROS.2011.6094879.

11. Fénelon, G., Soulas, T., De Langavant, L. C., Trinkler, I. & Bachoud-Lévi, A.-C. Feeling of presence in Parkinson's disease. *J Neurol Neurosurg Psychiatry* **82**, 1219–1224 (2011).

12. Bates, D., Mächler, M., Bolker, B. & Walker, S. Fitting Linear Mixed-Effects Models Using lme4. *Journal of Statistical Software* **67**, 1–48 (2015).

13. Kuznetsova, A., Brockhoff, P. B. & Christensen, R. H. B. lmerTest Package: Tests in Linear Mixed Effects Models. *Journal of Statistical Software* **82**, 1–26 (2017).

14. Ravina, B. *et al.* Diagnostic criteria for psychosis in Parkinson's disease: Report of an NINDS, NIMH Work Group. *Movement Disorders* **22**, 1061–1068 (2007).

15. Lenka, A., Pagonabarraga, J., Pal, P. K., Bejr-Kasem, H. & Kulisvesky, J. Minor hallucinations in Parkinson disease: A subtle symptom with major clinical implications. *Neurology* (2019) doi:10.1212/WNL.0000000000007913.

16. Ffytche, D. H. *et al.* The psychosis spectrum in Parkinson disease. *Nat Rev Neurol* **13**, 81–95 (2017).

17. Pagonabarraga, J. *et al.* Minor hallucinations occur in drug-naive Parkinson's disease patients, even from the premotor phase. *Mov. Disord.* **31**, 45–52 (2016).

18. Kataoka, H. & Ueno, S. Predictable Risk Factors for the Feeling of Presence in Patients with Parkinson's Disease. *Mov Disord Clin Pract* **2**, 407–412 (2015).

19. Hara, M. *et al.* A novel manipulation method of human body ownership using an fMRI-compatible master-slave system. *Journal of Neuroscience Methods* **235**, 25–34 (2014).

20. Oldfield, R. C. The assessment and analysis of handedness: The Edinburgh inventory. *Neuropsychologia* **9**, 97–113 (1971).

21. Blanke, O. *et al.* Neurological and Robot-Controlled Induction of an Apparition. *Curbio* **24**, 2681–2686 (2014).

22. Boes, A. D. *et al.* Network localization of neurological symptoms from focal brain lesions. *Brain* **138**, 3061–3075 (2015).

23. Boes, A. D. *et al.* Network localization of neurological symptoms from focal brain lesions. *Brain* **138**, 3061–3075 (2015).

24. Fox, M. D. Mapping Symptoms to Brain Networks with the Human Connectome. *New England Journal of Medicine* **379**, 2237–2245 (2018).

25. Nooner, K. B. *et al.* The NKI-Rockland Sample: A Model for Accelerating the Pace of Discovery Science in Psychiatry. *Front Neurosci* **6**, (2012).

26. Heydrich, L. & Blanke, O. Distinct illusory own-body perceptions caused by damage to posterior insula and extrastriate cortex. *Brain* **136**, 790–803 (2013).

27. Bejr-Kasem, H. *et al.* Disruption of the default mode network and its intrinsic functional connectivity underlies minor hallucinations in Parkinson's disease. *Mov. Disord.* **34**, 78–86 (2019).

28. Movement Disorder Society Task Force on Rating Scales for Parkinson's Disease. The Unified Parkinson's Disease Rating Scale (UPDRS): status and recommendations. *Mov. Disord.* **18**, 738–750 (2003).

29. Pagonabarraga, J. *et al.* Parkinson's disease-cognitive rating scale: a new cognitive scale specific for Parkinson's disease. *Mov. Disord.* **23**, 998–1005 (2008).
